# Local tree cover predicts mosquito species richness and disease vector presence in a tropical countryside landscape

**DOI:** 10.1101/2023.12.05.570170

**Authors:** Johannah E. Farner, Meghan Howard, Jeffrey R. Smith, Christopher B. Anderson, Erin A. Mordecai

## Abstract

**Context:** Land use change and deforestation drive both biodiversity loss and zoonotic disease transmission in tropical countrysides. For mosquito communities that can include disease vectors, forest loss has been linked to reduced biodiversity and increased vector presence. The spatial scales at which land use and tree cover shape mosquito communities present a knowledge gap relevant to both biodiversity and public health.

**Objectives:** We investigated the responses of mosquito species richness and *Aedes albopictus* disease vector presence to land use and to tree cover surrounding survey sites at different spatial scales. We also investigated species compositional turnover across land uses and along environmental gradients.

**Methods:** We paired a field survey of mosquito communities in agricultural, residential, and forested lands in rural southern Costa Rica with remotely sensed tree cover data. We compared mosquito richness and vector presence responses to tree cover measured across scales from 30m to 1000m, and across land uses. We analyzed compositional turnover between land uses and along environmental gradients of tree cover, temperature, elevation, and geographic distance.

**Results:** Tree cover was both positively correlated with mosquito species richness and negatively correlated with the presence of the common invasive dengue vector *Ae. albopictus* at small spatial scales of 90 – 250m. Land use predicted community composition and *Ae. albopictus* presence.

**Conclusions:** The results suggest that local tree cover preservation and expansion can support mosquito species richness and reduce disease vector presence. The identified spatial range at which tree cover shapes mosquito communities can inform the development of land management practices to protect both ecosystem and public health.

## Introduction

Humans have modified more than half of Earth’s land surface through activities such as deforestation, agricultural intensification and expansion, and urbanization (Hooke *et al*., 2012). Land use and land cover change fundamentally alter ecosystems, with strong impacts on biodiversity including species endangerment, loss, and invasion (e.g., Sala, 2000; Pereira *et al*., 2010; Maxwell *et al*., 2016; Giam, 2017). For communities containing species that transmit human diseases, the impacts of land use change on biodiversity can also impact human health. For example, the growing land area used for pineapple production in Costa Rica corresponds to an increase in highly suitable habitat for the malaria vector *Anopheles albimanus* (Rhodes *et al*., 2022). Developing land management solutions that benefit both biodiversity and public health is particularly important for countryside landscapes characterized by natural habitat remnants patchworked with human residential and agricultural infrastructure. These landscapes are globally dominant, are disproportionately impacted by land use change and zoonoses, and are critical to biodiversity conservation (Norris, 2008; Chazdon *et al*., 2009; Maxwell, 2016; Sokolow *et al*., 2022). Percent tree cover at small spatial scales of <100m has emerged as a reliable predictor of biodiversity for taxa in Latin American tropical countrysides including birds, non-flying mammals, and bats (Mendenhall *et al*., 2016), suggesting that local tree cover management holds promise as a practicable conservation tool. However, major knowledge gaps remain surrounding how local tree cover relates to invertebrate biodiversity, including for arthropod vectors of human diseases.

Mosquito communities (family Culicidae) are relevant to biodiversity and public health in countryside landscapes because they are sensitive to tree cover and habitat type, act as prey, predators, and detritivores in aquatic and terrestrial food webs (Addicott, 1974; Heard, 1994; Daugherty, Alto and Juliano, 2000; Poulin, Lefebvre and Paz, 2010), and can include species that are important vectors of diseases including malaria, dengue, chikungunya, Zika, yellow fever, West Nile fever, and arboviral encephalitis (Garmendia, Van Kruiningen and French, 2001; Lemon, 2008; LaPointe, Atkinson and Samuel, 2012; World Health Organization, 2021).

Previous studies show associations between forest conversion and low mosquito biodiversity, and suggest that high rates of land use change and active invasions by major vector species are together reshaping mosquito communities in ways that increase disease risk (Ferraguti *et al*., 2016; Meyer Steiger, Ritchie and Laurance, 2016; Burkett-Cadena and Vittor, 2018; Chaves *et al*., 2021). These patterns are likely shaped by abiotic and biotic conditions associated with tree cover and land use context that affect the presence and abundance of mosquito species that vary in their thermal niches, aquatic breeding habitat requirements, and preferred groups of vertebrates for blood meals (Laird, 1988; Gutman *et al*., 2004; Mordecai *et al*., 2019; Prevedello *et al*., 2019). Clarifying the spatial scales at which these landscape features shape mosquito communities is critical to understanding how tree cover management might balance biodiversity conservation, public health, and economic needs.

The countryside of Costa Rica is an ideal system in which to study relationships between land cover, mosquito community characteristics, and disease vector occurrence. This region, like neighboring Latin American countries, has a large burden of mosquito-borne diseases, including dengue virus, with two invasive *Aedes* spp. vectors potentially contributing to transmission (Rezza, 2012; Kraemer *et al*., 2015; WHO, 2021). *Aedes aegypti* is considered the primary dengue vector in Costa Rica, but the ongoing, patchily described invasion by the globally important vector species *Aedes albopictus* is potentially reshaping disease risk (Troyo *et al*., 2006; Calderón-Arguedas *et al*., 2010; Calderón-Arguedas *et al*., 2015; Rojas-Araya *et al*., 2017). Globally, *Ae. albopictus* and *Ae. aegypti* are predominantly associated with rural and urban human settlements, respectively, suggesting that *Ae. albopictus* and its responses to tree cover may play particularly important roles in countryside dengue transmission (Braks *et al*., 2003; Tsuda *et al*., 2006). Additionally, intensive long-term research in this system has characterized many links between landscape context, biodiversity, and ecosystem services (e.g., Daily *et al*., 2001; Ricketts *et al*., 2004; Karp *et al*., 2013; Frank *et al*., 2017; Hendershot *et al*., 2020; Langhans *et al*., 2022), but the corresponding links to mosquito biodiversity and disease vector presence remain undescribed.

Here, we combine field observations of mosquito communities in forested, agricultural, and residential settings in a rural area of southern Costa Rica with remotely-sensed land cover data in order to investigate mosquito community responses to forest cover and land use. We ask: How do mosquito community characteristics vary with tree cover measured at different spatial scales, among land use types, and along environmental gradients? We hypothesize that lower local tree cover and non-forest land uses are associated with lower mosquito species richness but higher presence of human-associated *Aedes* spp. vectors. By improving scientific understanding of how landscape context influences mosquito community assembly in Costa Rica, this study contributes to a more general understanding of links between land cover, biodiversity, and ecosystem services through the lens of entomological factors that influence disease risk.

## Methods

### Study area

This study was conducted in the cantons of Coto Brus, Corredores, and Golfito (8°43′14″N, 82°57′20″W), located in the southern Puntarenas region of Costa Rica along the border with Panama (Figure 1). The region ranges from coastal lowland tropical rainforest (0 m above sea level) to high elevation cloud forest (1500 m above sea level) and has distinct wet and dry seasons. The study area is predominately composed of rural communities surrounded by agriculture interspersed with forest patches, and also includes the protected Las Cruces forest reserve. In this area, dengue is endemic, *Ae. aegypti* is common, and *Ae. albopictus* has a growing presence (World Health Organization, 1994; Troyo *et al*., 2008, 2009; Rojas-Araya *et al*., 2017).

**Figure 1.**
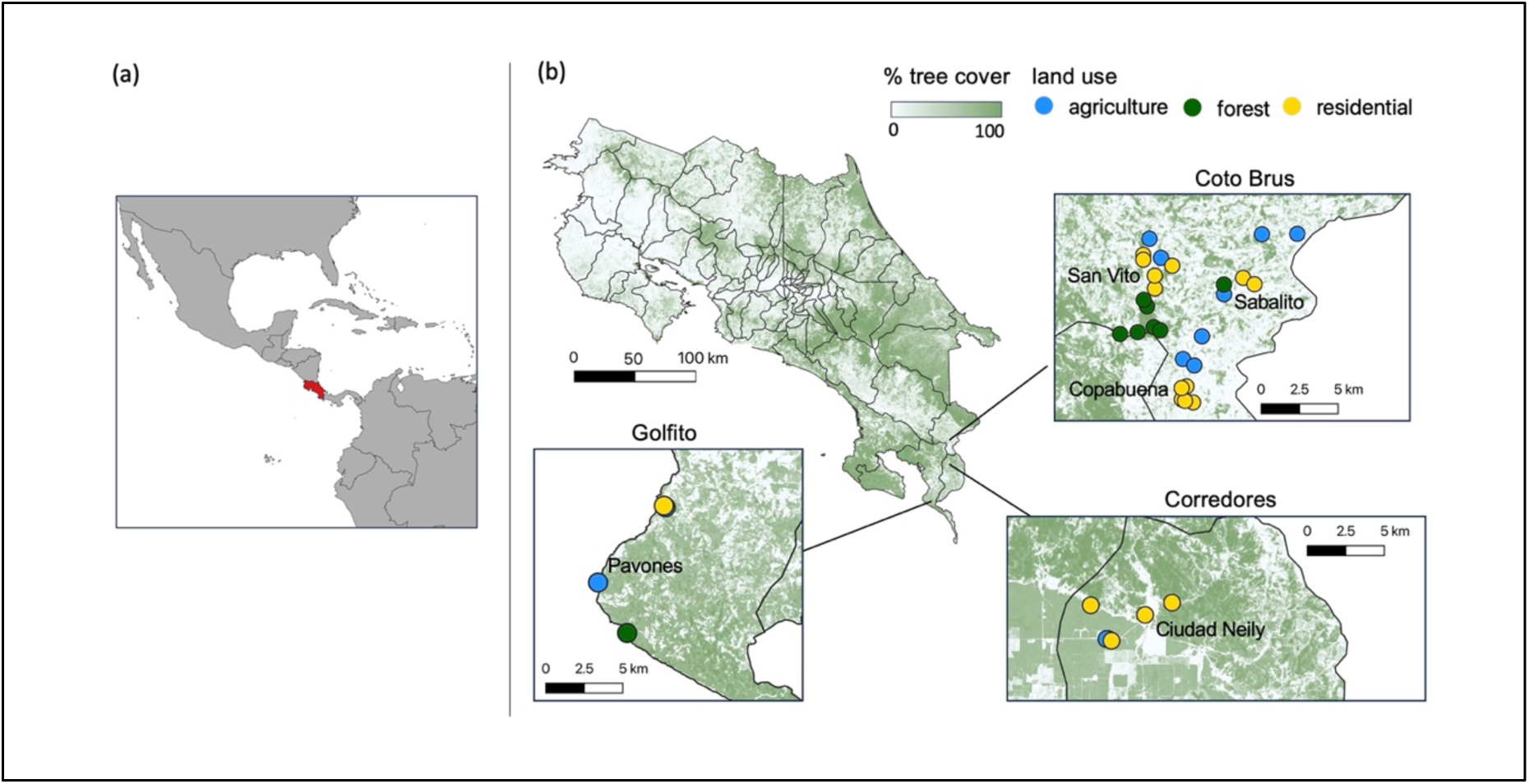
Tree cover and land use varied across study sites in Costa Rica. (A) Location of Costa Rica in Central America is highlighted in red. (B) Map of Costa Rica with canton boundaries delineated by lines, tree cover (remotely sensed at 30 m resolution) indicated in shades of green, and sampling sites indicated by colored points within the three insets. Insets shows study sites within Golfito, Corredores, and Coto Brus cantons. Blue, green, and yellow points denote agricultural, forest, and residential land uses, respectively. Text labels within insets denote districts.

### Study sites

With landowner permission, we accessed 37 sites representing three broad land use classes that were determined on-site by the survey team: residential (N = 17), agricultural (N = 12), and forest (N = 8) (Figure 1). In the Coto Brus canton (N = 12 residential sites, 8 agricultural, 7 forest), sites were in the districts San Vito, Sabalito, and Copabuena; in Corredores canton (N = 4 residential, 2 agricultural), sites were in Ciudad Neily district; in Golfito canton, (N = 1 residential, 2 agricultural, 1 forest), sites were in Pavones district. Residential site collections took place in the yards of residences in urban and peri-urban areas; agricultural site collections took place in the agricultural fields of coffee plantations, an oil palm plantation, a pine plantation, one mixed crop field, and one pasture; forest site trapping took place in primary and secondary forest edges, interiors, and fragments, on both protected and unprotected lands.

### Environmental variables

To quantify percent tree cover, we used a 30 m resolution map of tree cover in Costa Rica created by Echeverri et. al (2022) from multi-sensor satellite observations (2005 – 2017) and fine-scale tree cover maps (Figure 1). From this map, we calculated percent tree cover at different spatial scales surrounding each site using the R package “raster” (Hijmans *et al*., 2015). Specifically, we began by calculating percent tree cover within a radius of 30 m; we then increased this radius by increments of 10 m up to 200 m, and by increments of 50 m for radii between 200 m and 1000 m (following Mendenhall *et. al*, 2014). To account for mosquito interspecific variation in sensitivity to thermal conditions (Mordecai *et al.,* 2019), which could affect observed relationships between land cover and mosquito community characteristics along the 1500 m elevational gradient surveyed, we additionally extracted mean annual temperature data from 1970 – 2000 for each study site from the WorldClim 1 km^2^ resolution mean annual temperature dataset (Fick and Hijmans, 2017).

### Sample collection

We trapped mosquitoes during the rainy season, visiting all sites twice between June 19 and August 9, 2017, excepting the four sites in Pavones, which were trapped once (N = 70 trap nights). The interval between collections at a given site ranged from 8 to 36 days (mean = 22 days, SD = 8 days) (Table S1). In order to detect mosquito species with a variety of habitat and blood meal host preferences, we used a mix of trapping and manual collection methods (Thongsripong *et al*., 2013; Hoshi *et al*., 2014; Giordano *et al*., 2020; Romero-Vega *et al*., 2023). Specifically, at each site, we placed a total of four traps overnight for a 12-16 hour period within an area of 30 m in radius: one unlighted CDC trap baited with carbon dioxide produced by a mixture of Fleischmann’s Dry Active Yeast, household refined sugar, and water to mimic vertebrate respiration; one BG Sentinel baited with a BG-Lure and octanol to attract human-specialist mosquitoes; and two BG-GAT traps furnished with yellow sticky cards, corn oil, and a mixture of water and local leaf litter to attract gravid female mosquitoes. Trap locations within sites were chosen per BioGents recommendations, and square metal frames covered in large black plastic bags were placed over BG Sentinel and BG-GAT traps for protection from rainfall. To supplement the overnight trapping, during each trapping session, mosquito larvae were collected from breeding habitats and one of three trained members of the field team carried out 20 minutes of direct aspiration, over a standardized time and area.

### Mosquito identification

Four field team members trained to morphologically identify *Ae. albopictus* and *Ae. aegypti* picked out, sexed, and counted individuals of these species collected at each site. All morphological identifications were confirmed by the field team lead to ensure consistency. All other mosquitoes were counted, stored, and transported to Stanford University for molecular identification. We extracted and amplified DNA from the mitochondrial CO1 gene from pooled samples of mosquitoes from each trap night at each site (N = 70 pools) using MyTaq RedMix (Meridian Bioscience, Cincinnati, OH), following the protocol provided by the manufacturer.

Amplified DNA libraries were prepared for next-generation sequencing with Nextera (Illumina, Inc., San Diego, CA) and sequenced via Illumina MISEQ, with samples containing *Aedes albopictus* and *Culex tarsalis* DNA as positive controls. We removed primer sequences and paired forward and reverse reads from the sequencing data with the R package “dplyr”, and then used the R package “dada2” to filter and trim the DNA sequences to 473 bp, with a minimum overlap of 20 bases and a maximum of 5 expected errors (Callahan *et al*., 2016; Wickham *et al*., 2023). We estimated taxonomic placement for the sequenced mosquitoes by using the R packages “Biostrings” and “DECIPHER” to group DNA sequences into operational taxonomic units (OTUs, henceforth referred to as species) of 97% sequence similarity, and comparing representative sequences for each species to the BOLD and GenBank database records (Altschul *et al*., 1990; Ratnasingham and Hebert, 2007; Wright, 2016; Pagès *et al*., 2022). Species were identified based on top matches with sequence similarity ≥97% (Hardulak *et al*., 2020). When sequence similarity to the top match was < 97%, a higher level of taxonomic identification (e.g., genus) was assigned based on placement within a phylogenetic tree of the BOLD database sequences.

### Statistical analyses

We described mosquito communities in terms of species richness and species composition by combining presence data from the morphologically identified *Aedes* data and the sequencing data. To assess the completeness of species pool sampling for each land use type, we plotted species accumulation curves with the function “accumcomp” in the package BiodiversityR (Kindt and Coe, 2005). To quantify relationships between species richness and percent tree cover, we used generalized linear models (GLMs) with negative binomial error corrections for overdispersion and mean-centered independent variables. To assess the spatial scales at which tree cover best predicted species richness, we compared AIC values for GLMs that included percent tree cover surrounding each site calculated at radii ranging from 30 m to 1000 m. We used binomial logistic regression to analyze relationships between *Ae. albopictus* disease vector presence/absence and percent tree cover across spatial scales. For the 1000 m spatial scale where climate data were available, we additionally assessed the relative influence of mean annual temperature on species richness and *Aedes* vector presence with GLMs including mean annual temperature and its interaction with tree cover.

To compare species richness and *Aedes* vector presence between forest, agricultural, and residential land uses, we used Kruskal-Wallis tests with Bonferroni p-value adjustments to account for multiple comparisons.

To compare species composition among land use types and along environmental gradients, we first calculated the Jaccard coefficient of community similarity for each pair of sites for use in statistical tests and ordination. We then tested for differences in community similarity among land uses with permutational analysis of variation (PERMANOVA), first for all land use types, and then with pairwise adonis functions. Because PERMANOVA is sensitive to heterogeneity in dispersion among groups (Anderson and Walsh, 2013), we additionally tested whether dispersion differed among land use types using Tukey’s Honest Significant Differences method with betadisper() calculations of group average distances to the median. To visualize community similarity across land uses, we used non-metric multidimensional scaling (NMDS).

All the above analyses of compositional similarity among land uses were run using the R package “vegan” (Oksanen *et al*., 2013); for the pairwise PERMANOVA analyses, we used the R package “ecole” which provides wrapper functions for “vegan” (Smith, 2022). Finally, to quantify compositional turnover along environmental gradients of tree cover, mean temperature, elevation, and geographic distance, we used the R package “gdm” for generalized dissimilarity modeling (GDM), a form of nonlinear matrix regression that is robust to collinearity (Fitzpatrick *et al*., 2022). As above, we compared GDM models that incorporated tree cover at radii ranging from 30 – 1000 m to identify the spatial scale at which tree cover best explained compositional turnover.

All analyses were performed in R version 4.2.1. In addition to the R packages cited above, we used the packages “tidyverse” (Wickham *et al*., 2019), “cowplot” (Wilke *et al*., 2019), “MASS” (Ripley *et al*., 2013), “interactions” (Long, 2019), “gridExtra” (Auguie *et al*., 2017), and “reshape2” (Wickham, 2007) for data analysis and figure generation.

## Results

### Environmental variables

Across sites, tree cover ranged from 0% to 100% (mean = 27.9%, SD = 34.8%) at the smallest spatial scale we considered, a 30 m radius. Surrounding tree cover within a 1000 m radius, the largest spatial scale considered, ranged from 8.2% to 74.8% (mean = 33.4%, SD = 20.3%). The lowest and highest site elevations were 12 m and 1451 m above sea level, respectively (mean = 776 m, SD = 473.2 m). Mean annual temperature ranged from 18.8 to 26.4 °C (mean = 22.6 °C, SD = 2.3 °C).

### Mosquito collections

A total of 1,283 mosquitoes representing 48 species in 13 genera were collected from 37 sites (Figure 2, Table S2). The number of mosquitoes collected at a site ranged from one to 244 (mean = 35, SD = 64) (Table S1). Of these, 99 individuals from 14 residential and five agricultural sites were morphologically identified as *Ae. albopictus*, and five total *Ae. aegypti* individuals were identified from one forest, one agricultural, and one residential site. *Ae. albopictus* DNA was detected in the pooled samples of molecularly identified mosquitoes from seven of the 19 sites where *Ae. albopictus* individuals were also morphologically identified, and no sites without morphologically identified specimens. The five most common species were *Ae. albopictus*, *Culex quinquefasciatus*, *Cx. nigripalpus*, *Wyeomyia adelpha/Guatemala,* and *Limatus durhamii* (Figure 2, Table S3).

**Figure 2.**
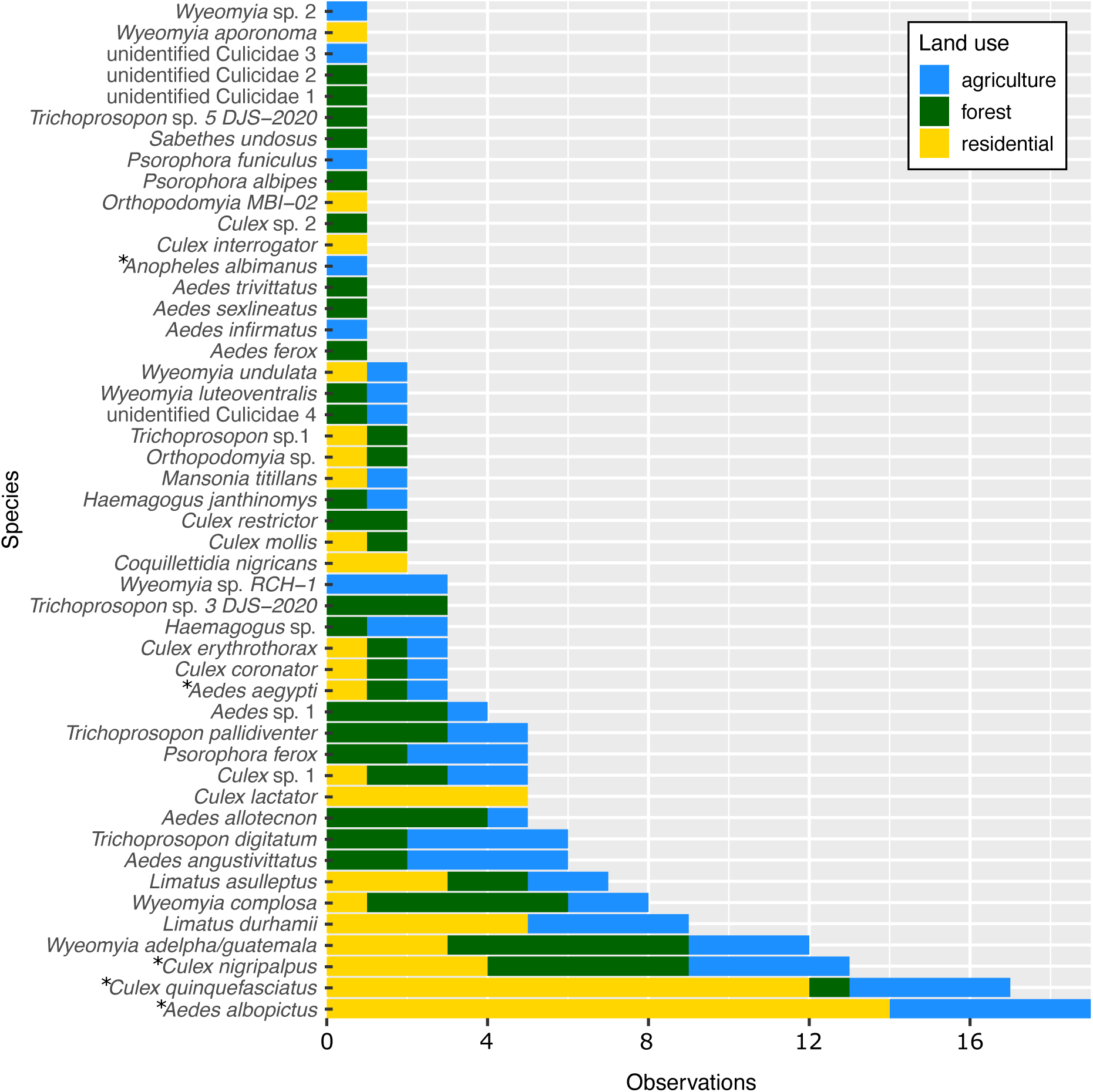
The commonness of 48 observed mosquito species varied both overall and among land use types. Blue bars show the number of agricultural sites in which each species was present, green bars show forest sites, and yellow bars show residential sites. Asterisks indicate species known to vector human diseases.

### Species richness

Site-level species richness ranged from one to 19 (mean = 4.89, SD = 4.1) (Table S2). Overall species counts for forest, agricultural, and residential land uses were 33, 29, and 21, respectively. Ten species (21%) were observed in all three land uses. Nineteen species (40%) were shared among forest and agricultural land uses, 13 species (27%) were shared among agricultural and residential land uses, and twelve species (25%) were shared among forest and residential land uses (Figure 2, Table S3). Eleven species (23%) were found only in forested settings, six species (13%) were found only in agricultural settings, and six species (13%) were found only in residential settings (Figure 2, Table S3). Two species, *Ae. albopictus* and *Culex quinquefasciatus*, were common (observed at >50% of sites) in residential settings, no species were common in agricultural settings, and three species—*Culex nigripalpus*, *Wyeomyia complosa*, and *Wyeomyia adelpha/guatemala*—were common in forested settings (Table S1). Species accumulation curves indicate that more species would have been observed in each land use class with additional sampling, but at a lower rate in residential compared to forest and agricultural land uses (Figure S1).

### Disease vectors

At least five of the mosquito species observed are known vectors of human diseases. Three of these—the dengue and chikungunya virus vector *Ae. albopictus* (present at 19 sites) and the St. Louis Encephalitis virus vectors *Cx. quinquefasciatus* (present at 17 sites) and *Cx. nigripalpus* (present at 13 sites)—were the three most frequently observed species (Reisen, 2003; Simmons *et al*., 2012). Rarely observed vector species included the dengue, chikungunya, yellow fever, and Zika virus vector *Ae. aegypti* (present at three sites spanning all three land use types) and the malaria vector *Anopheles albimanus* (present at one agricultural site) (Zimmerman, 1992; Simmons *et al*., 2012). In contrast to *Ae. aegypti, Cx. nigripalpus,* and *Cx. quinquefasciatus*, which were observed in all land use types, *Ae. albopictus* was observed only in residential and agricultural settings associated with intensive human modification.

### Model results

Mosquito species richness was explained by tree cover, but not by land use type. Comparisons of GLMs using tree cover calculated for radii ranging between 30 m and 1000 m surrounding each site indicated that species richness was positively correlated with tree cover at radii between 90 m and 650 m, and tree cover at a 250 m radius had the largest effect size (estimated effect = 1.40 x 10^-2^, SE = 5.00 x 10^-3^, z-value = 2.8, p-value = 5.08 x 10^-3^) (Figure 3a, Table S4). At the 1000 m spatial scale where both tree cover and climate data were available, the interaction between tree cover and mean annual temperature had a significant effect on species richness (estimated effect = −9.48 x 10^-3^, SE = 3.48 x 10^-3^, z-value = −2.72, p-value = 6.48 x 10^-3^ (Table S5). Specifically, at high temperatures, species richness was low even when tree cover was high (Figure S2). By contrast, Kruskal-Wallis test results indicated that species richness did not differ significantly among forested, agricultural, and residential sites (chi-squared = 2.84, df = 2, p-value = 0.24) (Figure 3b). Notably, the highest species richness was observed at a site in the Coto Brus forest reserve (Figure 3b).

**Figure 3.**
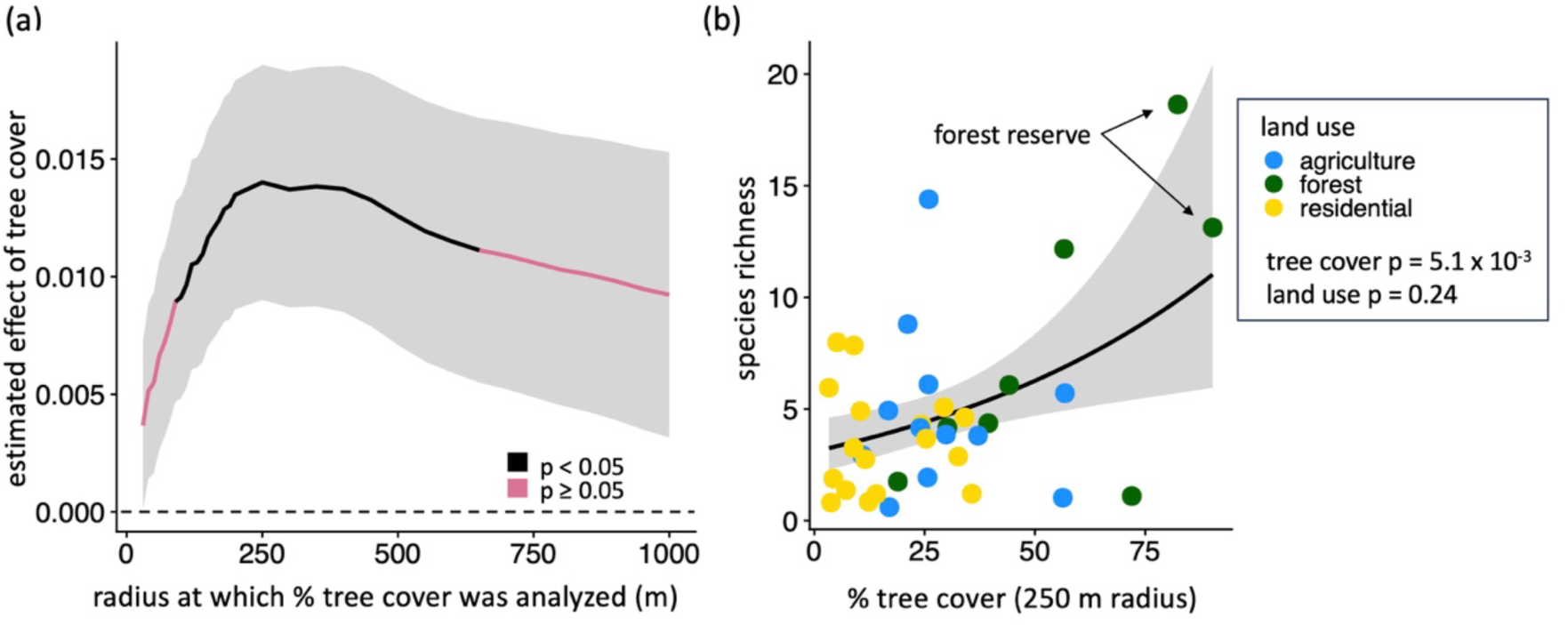
Species richness is correlated with tree cover surrounding survey sites for radii between 90 m and 650. **m.** *(a)* The estimated effect of surrounding tree cover calculated across spatial scales on species richness. Radii where the relationship between tree cover and species richness is significant (p < 0.05) are shown in black; others are shown in pink. Values above the dashed line are positive. (b) Species richness increases with percent tree cover at a 250 m radius: the scale identified as having the strongest effect. Land use (colored points) and species richness are not significantly correlated. The site with the highest species richness and the site with the highest surrounding tree cover were both located in the Las Cruces forest reserve (arrows). In both panels, gray shading shows +/- 1 SE.

### Relationships of Ae. albopictus presence to tree cover and land use

From the *Aedes* disease vector survey, we present results only for *Ae. albopictus* because observations of *Ae. aegypti* were insufficient for statistical analysis. Both tree cover and land use type predicted *Ae. albopictus* presence. Comparisons of GLMs using tree cover calculated for radii ranging between 30 m and 1000 m surrounding each site indicated that *Ae. albopictus* presence was negatively correlated with tree cover at radii between 30 m and 250 m, and was best explained by tree cover at a 110 m radius (estimated effect = −4.46 x 10^-2^, SE = 1.83 x 10^-2^, z-value = −2.44, p-value = 1.47 x 10^-2^) (Figure 4a, Table S6). At the 1000 m spatial scale where we additionally assessed the influence of climate, *Ae. albopictus* presence was negatively correlated with tree cover and positively correlated with temperature (tree cover estimated effect = −8.34 x 10^-2^, SE = 3.61 x 10^-2^, z-value = −2.31, p-value = 2.08 x 10^-2^; mean annual temperature estimated effect = 9.36 x 10^-1^, SE = 4.2 x 10^-1^, z-value = 2.23, p-value = 2.5 x 10^-2^) (Figure S3, Table S7). Land use type also predicted *Ae. albopictus* presence (Kruskal-Wallis chi-squared = 15.02, p-value = 5.48 x 10^-4^). Specifically, *Ae. albopictus* was significantly more likely to be observed in residential settings (present at 14/17 sites) than in forested settings (present at 0/8 sites) (Kruskal-Wallace chi-squared = 14.37, Bonferroni-adjusted p-value = 4.49 x 10^-4^), and its presence in agricultural settings (present at 5/12 sites) did not differ significantly compared to either residential (Kruskal-Wallace chi-squared = 4.98, adjusted p-value = 7.8x 10^-2^) or forested (Kruskal-Wallace chi-squared = 4.22, adjusted p-value = 1.20 x 10^-1^) settings. Eighteen of the 19 sites where *Ae. albopictus* was present were surrounded by less than 35% tree cover within a 110 m radius. The only site within a pine plantation was a clear outlier, where *Ae. albopictus* was present under 75% tree cover (Figure 4B).

**Figure 4.**
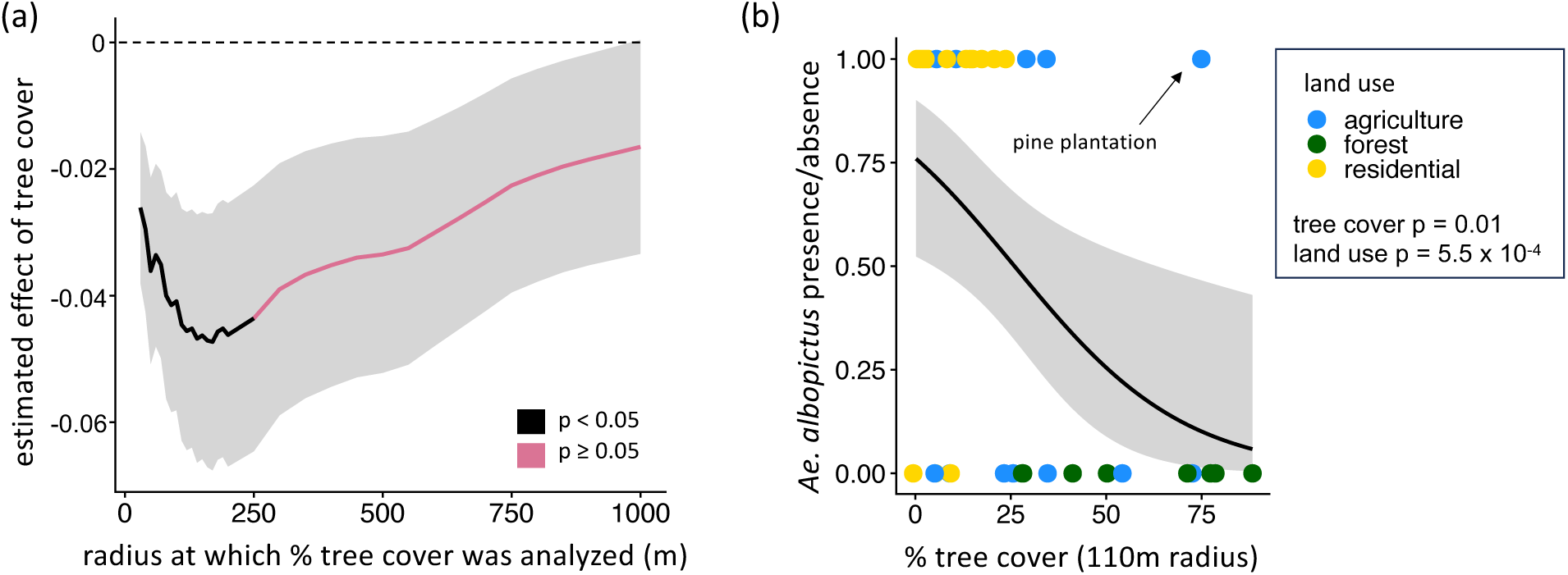
*Aedes albopictus* is most likely to be found at low tree cover levels and in residential settings. *(a)* The estimated effects of tree cover across spatial scales on Ae. albopictus presence, where black lines indicate statistically significant (p < 0.05) effects and pink lines indicate non-significant (p > 0.05) relationships. Values below the dashed line are negative. (b) Land use type (colored points), percent tree cover, and Ae. albopictus presence/absence at the highest-significance 110 m radius. In both panels, gray shading shows +/- 1 SE.

### Species composition across land uses

In contrast to species richness, community composition and dispersion were predicted by land use type. PERMANOVA results comparing all three land uses showed that land use significantly affected community composition (sum of squares = 2.24, R^2^ = 0.16, F-value = 3.24, p-value = 0.001), and pairwise PERMANOVAs showed that agricultural and forest mosquito communities differed significantly from residential communities (Table 1). Wider dispersion among agricultural compared to residential mosquito communities (average distance to the median: agriculture = 0.618, forest = 0.572, residential = 0.496; Tukey test adjusted p-values: residential – agricultural = 0.0104, residential – forested = 0.218, forested – agricultural = 0.604) likely contributed to the community dissimilarity detected between these land uses (Anderson and Walsh, 2013). Supporting these statistical results, NMDS visualization of communities grouped by land use type highlights that agricultural communities bridge distinct forest and residential communities (NMDS stress = 0.12) (Figure 5). The wider variation among agricultural sites is also evident from the species observation table: no single species was observed at more than 1/3 of all agricultural sites, whereas *Ae. albopictus* and Unidentified Culicidae 1 were both observed at >70% of residential sites, and *Culex nigripalpus*, *Wyeomyia adelpha/guatemala*, and *Wyeomyia complosa* were each observed at >60% of forested sites (Tables S2, S3). On the NMDS plot, forest land use sites falling far outside of the 95% CI included the single forest fragment and two of four forest edges surveyed (Figure S4).

**Figure 5.**
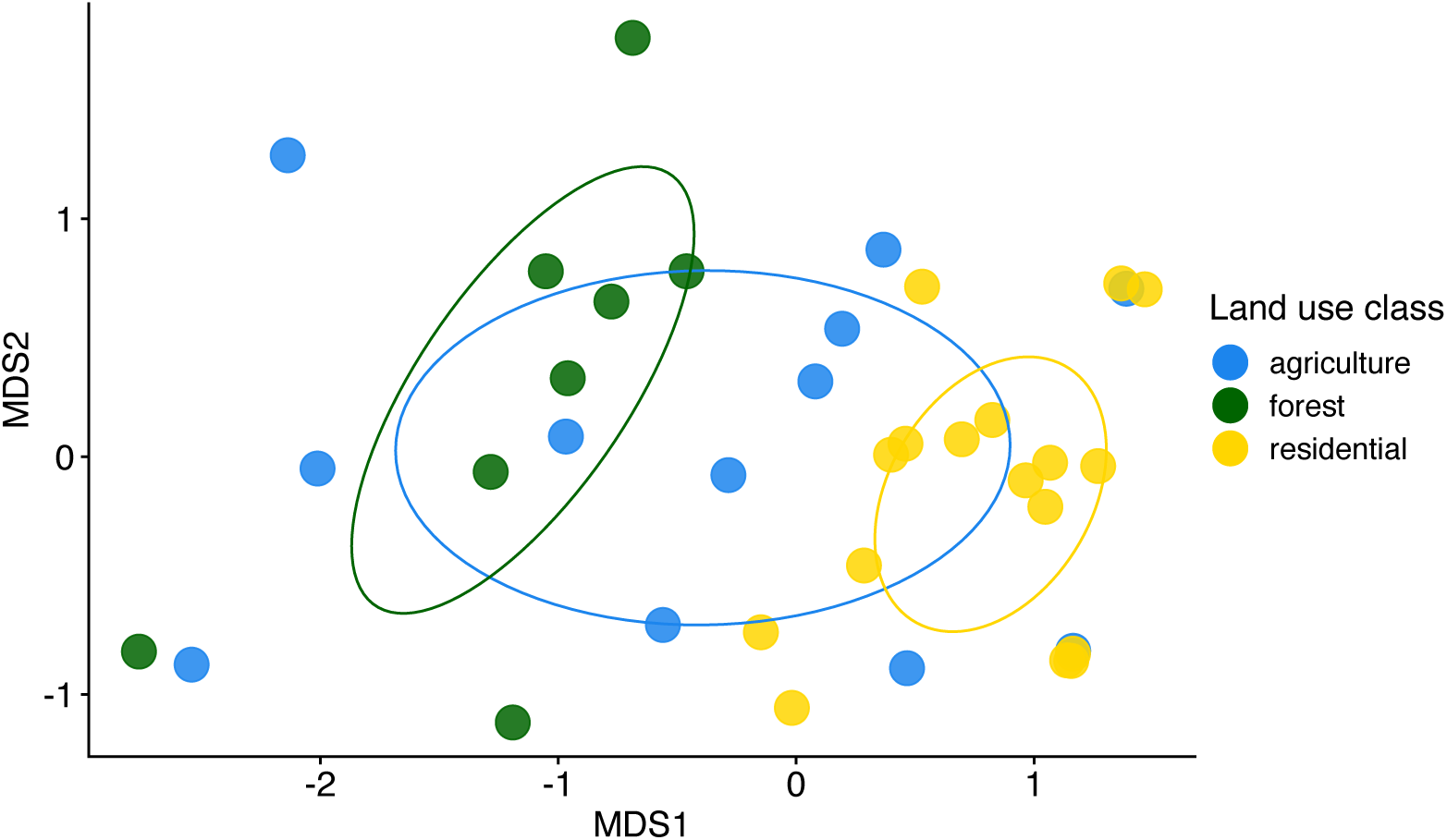
Distinct mosquito communities were observed in forest and residential land uses, while communities in agricultural settings overlapped with both other land use types. NMDS ordination visualization groups sites by community similarity, with colors indicating land use types (blue: agriculture, green: forest, yellow: residential). Each point represents the community at one study site, and the distance between points is smaller for more similar communities. Ellipses show 95% confidence intervals for ordination of agriculture, forest, and residential mosquito communities.

**Table 1.** PERMANOVA results for community composition compared among land use types. Asterisks indicate p < 0.05.

| Land use pair | Sum of squares | F-Value | R<sup>2</sup> | Bonferroni-adjusted p-value |
| --- | --- | --- | --- | --- |
| agriculture vs. forest | 0.645 | 1.59 | 0.0813 | 0.117 |
| agriculture vs. residential | 0.846 | 2.52 | 0.0853 | 0.018 * |
| forest vs. residential | 1.82 | 5.87 | 0.203 | 0.003 * |

Agricultural land use outliers clustering near the residential group included both oil palm plantations surveyed and one coffee plantation, and those clustering nearer the forest group included three coffee plantations (Figure S4). Each site in the geographically distinct Pavones district that was sampled only once fell outside of the 95% CIs for its land use type (Figure S4).

### Species turnover along environmental gradients

Finally, generalized dissimilarity modeling (GDM) indicated that environmental gradients explained little of the species turnover among sites. Among the spatial scales for which tree cover was calculated, the model using the 130m radius explained the highest amount of species turnover among sites (Table S8). The model that included mean annual temperature, geographic distance, and tree cover at the 130m radius explained 7% of species turnover among sites. Elevation showed no relationship with species turnover. Whereas increasing tree cover was associated with a consistent increase in community turnover, increasing temperature was associated with a steep increase in community turnover up to a plateau around 22 °C, and increasing geographic distance was associated with comparatively limited turnover (Figure 6).

**Figure 6.**
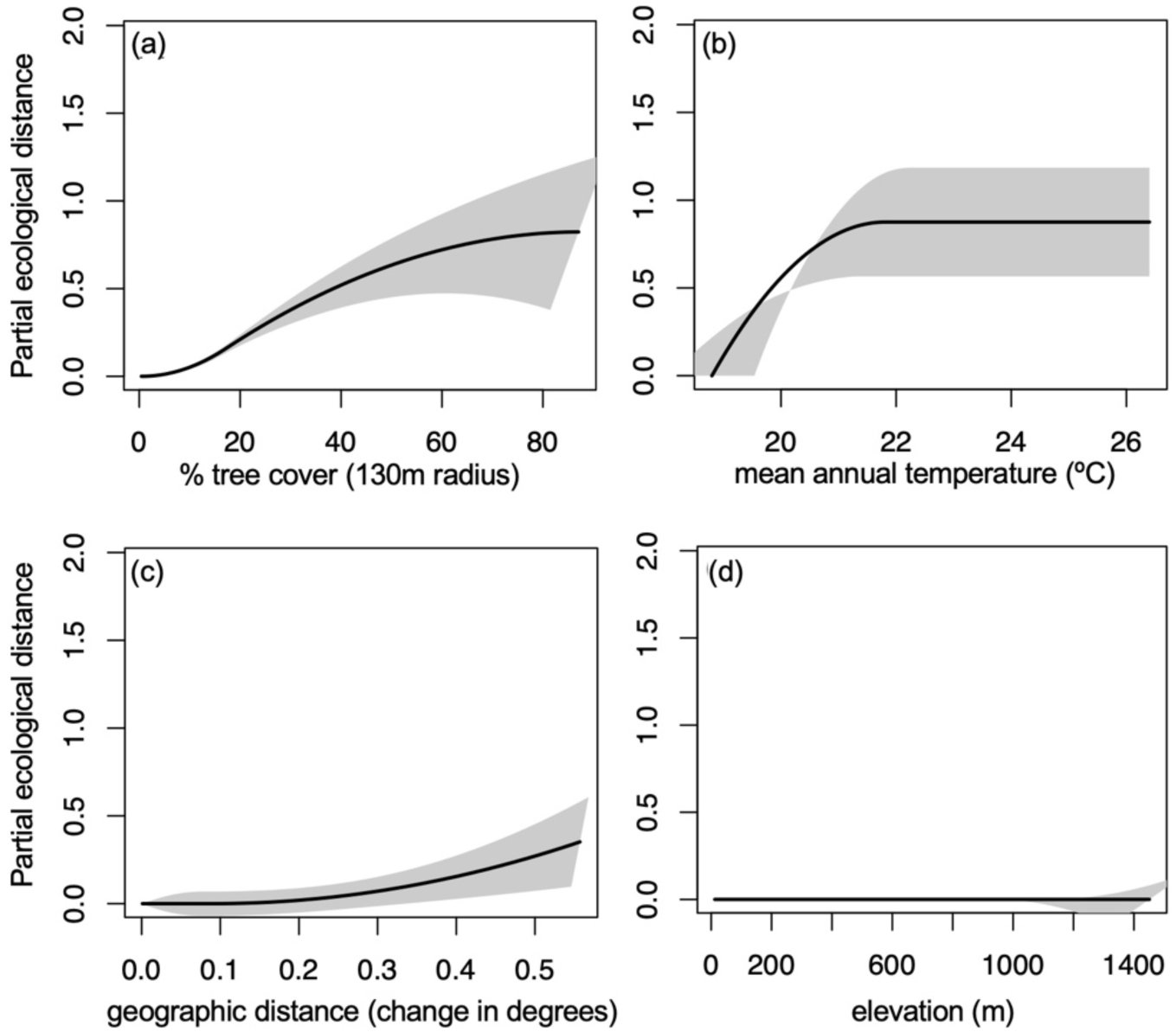
Generalized dissimilarity modelling (GDM) of community dissimilarity indicated that (a) percent tree cover and (b) mean annual temperature explained 7% of deviation from the null. (c) geographic distance contributed minimally and (d) elevation did not significantly contribute to community turnover. In each panel, the x-axis shows the environmental gradient and the y-axis shows the amount of compositional turnover, measured as partial ecological distance. The maximum height the spline reaches on the y-axis indicates the total amount of compositional turnover the gradient is associated with, and the slope shows how the rate of compositional turnover varies along the environmental gradient. The difference in height between any two points along the I-spline corresponds to the modeled contribution of that predictor variable to the difference between those points. Grey shading shows +/- 1 standard deviation when 70% of sites are sampled 10 times.

## Discussion

We found that local tree cover, but not land use (residential, agricultural, or forest), predicted mosquito species richness, suggesting that more diverse communities occur at higher tree cover. By contrast, community composition was more predictable for forested and residential land uses, and more variable among agricultural sites, but environmental gradients of tree cover, climate, and geographic distance explained only 7% of species turnover among sites. *Ae. albopictus* presence varied significantly with both tree cover and land use, but in the opposite direction from mosquito richness: increasing with lower tree cover and in residential compared to forested sites (with intermediate probability in agricultural sites). Overall, our results add to support from both mosquitoes and other taxa that natural and semi-natural habitat, including tree cover, sustains substantial biodiversity and ecosystem services—here, in the form of protection against mosquitoes that vector human disease (Millennium Ecosystem Assessment, 2005; Jose, 2009; Mendenhall *et al*., 2014, 2016; Frank *et al*., 2017; Barrios *et al*., 2018; Burkett-Cadena and Vittor, 2018; Frishkoff *et al.,* 2019; Langhans *et al*., 2022; Perrin *et al*., 2023).

### Mosquito richness and species composition

The spatial scales at which tree cover predicted mosquito species richness were small (90-650 m), and comparable to previous findings for other taxa in the same study area, highlighting the disproportionately positive impact of small patches of trees on biodiversity (Mendenhall *et al*. 2014, 2016; Frank *et al*., 2017). The radius at which tree cover best predicted species richness was 250 m; by comparison, biodiversity was correlated with tree cover at small spatial scales for non-flying mammals (70 m), bats (50 - 60 m), birds (30 m), reptiles (50 m), and amphibians (80 m) in the same region of Costa Rica (Mendenhall *et al.,* 2014, 2016; Frank *et al.,* 2017). Although the radius at which mosquito species richness responded most strongly to tree cover was larger compared to previously studied taxa, the spatial scale remained local, and significant effects of tree cover were found at radii as small as 90 m. Our observation of a positive relationship between species richness and tree cover aligns with those of many other studies of mosquito diversity along land cover gradients in locations including Latin America, Asia, and Europe (e.g., Johnson, Gómez and Pinedo-Vasquez, 2008; Thongsripong *et al*., 2013; Ferraguti *et al*., 2016; Chaverri *et al*., 2018), and our analysis of the distance around sampling sites at which tree cover shapes mosquito communities contributes to clarifying the spatial scales at which mosquitoes respond to landscape features.

Our observation that mosquito community composition was distinct among different land uses is consistent with patterns observed both for other taxa in this system, and for mosquitoes in other regions (Mendenhall *et al*., 2014, 2016; Meyer Steiger, Ritchie and Laurance, 2016; da Silva Pessoa Vieira *et al*., 2022). In agricultural settings, relatively high species richness and community similarity with forested settings support the argument that farmlands can contribute substantially to biodiversity (e.g., Norris, 2008). However, the high proportion of species unique to forest habitats and the high species richness observed inside the large Las Cruces forest reserve also reaffirm the importance of forests and protected areas as refugia for biodiversity (Coetzee, Gaston and Chown, 2014; Mendenhall *et al*., 2016). Additionally, the compositional variability among agricultural settings and the close community resemblance between some agricultural and residential sites indicate a need for additional research on how mosquito communities respond to specific land uses, crop assemblages, or management practices that can result in similar levels of tree cover. For example, organic farming methods are associated with higher arthropod diversity globally compared to conventional methods, and Kenyan ricelands that rely on natural rather than artificial irrigation have higher mosquito species richness (Muturi *et al*., 2006; Lichtenberg *et al*., 2017). Costa Rican croplands that are less intensively farmed support greater bird species richness, a pattern that may also hold for mosquitoes (Hendershot *et al*., 2020).

Environmental gradients of tree cover and temperature shaped species turnover, but explained only 7% of variance in community composition, suggesting that additional habitat characteristics may play important roles in determining species composition. Such factors might include local microclimates, differences among types of tree cover (e.g., agricultural types, primary versus secondary forest), and/or the presence of vertebrate hosts preferred by different mosquito species. Differences in species abundances and community evenness, which were not quantified here, might also respond more strongly to gradual environmental change than the identities of the species present. Additionally, undersampling of communities may have contributed to this result by limiting the repeatability of community composition observed at environmentally similar sites. However, our result that land use predicts species composition, while land cover predicts species richness, aligns with patterns of abundance-based Dipteran and Culicid diversity observed in the tropical Australian countryside (Smith and Mayfield, 2015; Meyer Steiger, Ritchie and Laurance, 2016).

### Disease vectors

The most frequently observed disease vector, *Ae. albopictus*, was more likely to be observed in sites with lower surrounding tree cover and agricultural or residential land uses, suggesting that rural landscapes with more forest and tree cover may be more resistant to invasion by this species. These observations align with this species’ well-established preferences for taking blood meals from humans and livestock (Niebylski *et al*., 1994; Richards *et al*., 2006), and its association with rural, agricultural, suburban, and/or deforested settings in the Americas, Asia, and Africa (Gilotra, Rozeboom and Bhattacharya, 1967; Braks *et al*., 2003; Young *et al*., 2017; Câmara *et al*., 2020; Canelas *et al*., 2023). The 30 – 250 m radii at which tree cover negatively affected *Ae. albopictus* presence fell within the 90 – 650 m range at which tree cover positively affected species richness, suggesting promise for local tree cover management as a means of supporting both public health and biodiversity conservation. Protection against mosquito disease vectors conferred by tree cover may extend beyond *Ae. albopictus* to include at least 16 other significant vectors of human diseases that are favored by deforestation, including *Ae. aegypti,* multiple Anopheline malaria vectors, and the Amazonian malaria vector *Nyssorhynchus darlingi* (Burkett-Cadena and Vittor, 2018; Chaves *et al*., 2021).

Our finding that *Ae. albopictus* was associated with, but inconsistently observed in, agricultural settings (present in 33% of agriculture sites) reinforces that agricultural lands have the potential to either harbor or resist invasive species, and suggests vector associations with agricultural subtypes as a key future research direction. In our survey, factors that differentiated the high tree-cover pine plantation and two of the six surveyed coffee plantations as suitable habitat for *Ae. albopictus* are of particular interest. For example, *Ae. albopictus* larvae were found in banana leaves and stumps in a shade-grown coffee plantation. Understanding vector responses to agriculture is particularly important because this land use is the most likely candidate for local tree cover management in Costa Rica due to its spatial extent, the established Payment for Ecosystem Services program for incentivizing landowner forest retention and tree planting, and a previous finding that urban tree cover is correlated with dengue incidence in this region, while forest cover at the district level is negatively associated with dengue hospitalizations and outbreaks (Sánchez-Azofeifa *et al*., 2007; Troyo *et al*., 2009; Piaggio, 2024).

The findings of this study are subject to several limitations resulting from incomplete sampling of the mosquito community. Extensive sampling is required to fully capture the high arthropod biodiversity present in tropical areas (Coddington *et al*. 2009), such that unsaturated species accumulation curves are common in these systems (e.g., Novotný and Basset, 2003; Responte and Nuneza, 2016; Thormann *et al*., 2016; Romero-Vega *et al*., 2023; Kirmse, 2024). Many more mosquito species may have been observed with increased collection time at each site, particularly with additional coverage to include the dry season, as highlighted by seasonal mosquito species richness and abundances observed in Costa Rica by Romero-Vega *et al*. (2023). The sequencing approach we took to identify species other than *Ae. albopictus* and *Ae. aegypti* allowed us to efficiently identify specimens compared to a traditional morphological approach, but was limited by sequencing success and the availability of database entries for comparison. Additionally, the pooling of sequenced mosquitoes by site and trap night prevented measurements of species abundances and evenness, which are likely sensitive to tree cover and are key indicators of community structure and functioning, including the potential for vector species to transmit diseases (Franklinos *et al*., 2019). Despite these limitations, the number of mosquitoes collected and species richness we observed are on par with other mosquito studies in Costa Rica (Calderón-Arguedas *et al*., 2008; Burkett-Cadena *et al.,* 2013; Chaverri *et al*., 2018; Romero-Vega *et al*., 2023), and the statistically significant environmental responses of mosquito community characteristics were consistent with hypotheses grounded in the findings of other studies with much larger sample sizes (e.g., Braks *et al*., 2003; Johnson, Gómez and Pinedo-Vasquez, 2008; Chaves *et al*., 2021; da Silva Pessoa Vieira *et al*., 2022). We hypothesize that additional sampling in this system might uncover more species in all sites and land use types but also (1) significantly lower species richness in residential compared to forest and agricultural land uses, based on the comparatively shallow slope of the residential species accumulation curve; (2) a stronger signal of species turnover along environmental gradients, based on the large uncertainty in the GDM results; and (3) rare occurrences of *Ae. albopictus* in forest edge habitats, based on the association of this species with the urban-forest interface in Brazil (Pereira dos Santos *et al*., 2018; Hendy *et al*., 2023).

To improve understanding of how local tree cover shapes mosquito biodiversity and public health risk, future studies should aim to capture species richness and abundance along tree cover gradients and across seasons (Calderón-Arguedas *et al*., 2008; Troyo *et al*., 2008; Romero-Vega *et al*., 2023); assess effects of other factors in habitats that have similar tree cover but differ in aspects such as crop type or tree cover geometry; and use reforestation efforts or experimental tree cover additions at relevant spatial scales to test mosquito community responses. In the study region, tree cover responses of *Ae. aegypti* and *An. albimanus* require clarification, as we observed these regionally important vectors too rarely for statistical analysis; in particular, the canonically widespread and urban vector *Ae. aegypti* was unexpectedly rare and generalist, occurring in one site of each of the three land use types (Troyo *et al*., 2006; Cáceres *et al*., 2012). Additionally, the extent to which local tree cover effects on vector presence shape the potential for disease transmission should be tested with human pathogen surveillance in field-captured mosquitoes from different environments, and by comparison of disease case data to mosquito community data. In addition to dengue, Zika, and malaria, St. Louis Encephalitis Virus should be considered for surveillance in rural areas where humans and animals live in close proximity, because two potential vectors—*Cx. quinquefasciatus* and *Cx. nigripalpus—*were common, and this unmonitored disease is already widespread among both domesticated and wild animal hosts in Costa Rica (Medlin *et al*., 2016; Chaves *et al*., 2021; Piche-Ovares *et al*., 2023).

## Conclusions

Overall, our findings follow patterns observed repeatedly across the globe associating tree cover with higher mosquito species richness and lower disease vector presence. We showed that tree cover both increases mosquito species richness and decreases *Ae. albopictus* presence at small spatial scales of 90 – 250m. We also found that at larger spatial scales of 1000m, warm mean annual temperatures increase habitat suitability for *Ae. albopictus* and limit tree cover contributions to species richness, but note that other factors that covary with climate across the study region may contribute to this result. Although the specific mechanisms and characteristics by which tree cover inhibits disease vectors remain unclear, the alignment of local tree cover effects on mosquito communities with other benefits for biodiversity and ecosystem services adds support to the idea that countryside landscapes can be managed to foster both human and ecosystem health.

## Acknowledgements

We thank the many landowners who generously provided access to research sites; Henry Sandi Amador, Esteban Calderon, and Erick Barrantes of the Organization for Tropical Studies for field assistance; Jamieson O’Marr for field and laboratory assistance; Gretchen Daily for advice on the project design; and Mordecai lab members for laboratory assistance and helpful feedback throughout all stages of the project. This work was funded by the National Institutes of Health (R35GM133439, R01AI168097, R01AI102918), the National Science Foundation (DEB3322011147, with Fogarty International Center), the Stanford Center for Innovation in Global Health, King Center on Global Development, Woods Institute for the Environment, the Ward Wilson Woods Jr Environmental Study Fund, the Lewis and Clark Fund from the American Philosophical Society, and the Bing-Mooney Fellowship.

## Permits

Field collections and sample exports were carried out under general insect collecting permit R-036-2017-OT-CONAGEBIO, Costa Rica export permit 434-DGVS-2016, USDA veterinary import permit 134109, and Health and Human Services permit 2017-08-079.

## Author contributions

All authors contributed to the study conception and design. MH, JS, and CA collected the data. MH and JF prepared the data, and JF analyzed the data. The first draft of the manuscript was written by JF, and all authors contributed to subsequent versions of the manuscript. All authors read and approved the final manuscript.

The authors have no competing interests to declare.

## Data availability

The data presented in this manuscript are available in the DRYAD repository at https://doi.org/10.5061/dryad.p8cz8w9xg.

## References

Addicott, J.F. (1974) ‘Predation and prey community structure: An experimental study of the effect of mosquito larvae on the protozoan communities of pitcher plants’, Ecology, 55(3), pp. 475–492. Available at: 10.2307/1935141.

Altschul, S.F., Gish, W., Miller, W., Myers, E.W., and Lipman, D.J. (1990) ‘Basic local alignment search tool’, Journal of Molecular Biology, 215(3), pp. 403–410. Available at: 10.1016/S0022-2836(05)80360-2.

Anderson, M.J. and Walsh, D.C.I. (2013) ‘PERMANOVA, ANOSIM, and the Mantel test in the face of heterogeneous dispersions: What null hypothesis are you testing?’, Ecological Monographs, 83(4), pp. 557–574. Available at: 10.1890/12-2010.1.

Auguie, B., Antonov, A. and Auguie, M.B., 2017. ‘Package ‘gridExtra’’, Miscellaneous Functions for “Grid” Graphics.

Barrios, E., Valencia, V., Jonsson, M., Brauman, A., Hairiah, K., Mortimer, P.E., and Okubo, S. (2018) ‘Contribution of trees to the conservation of biodiversity and ecosystem services in agricultural landscapes’, *International Journal of Biodiversity Science*, Ecosystem Services & Management, 14(1), pp. 1–16. Available at: 10.1080/21513732.2017.1399167.

Braks, M.A.H., Honório, N.A., Lourenço-De-Oliveira, R., Juliano, S.A., and Lounibos, L.P. (2003) ‘Convergent habitat segregation of *Aedes aegypti* and *Aedes albopictus* (Diptera: Culicidae) in Southeastern Brazil and Florida’, Journal of Medical Entomology, 40(6), pp. 785– 794. Available at: 10.1603/0022-2585-40.6.785.

Burkett-Cadena, N., Graham, S.P. and Giovanetto, L.A. (2013) ‘Resting environments of some Costa Rican mosquitoes’, Journal of Vector Ecology, 38(1), pp. 12–19. Available at: 10.1111/j.1948-7134.2013.12004.x.

Burkett-Cadena, N.D. and Vittor, A.Y. (2018) ‘Deforestation and vector-borne disease: Forest conversion favors important mosquito vectors of human pathogens’, Basic and Applied Ecology, 26, pp. 101–110. Available at: 10.1016/j.baae.2017.09.012.

Cáceres, L., Rovira, J., Torres, R., García, A., Calzada, J., and De La Cruz, M. (2012) ‘Characterization of *Plasmodium vivax* malaria transmission at the border of Panamá and Costa Rica’, Biomedica, 32(4), pp. 557–569. Available at: 10.1590/S0120-41572012000400011

Calderón-Arguedas, O., Troyo, A., Solano, M.E., Avendaño, A., and Beierg, J.C. (2008) ‘Urban mosquito species (Diptera: Culicidae) of dengue endemic communities in the Greater Puntarenas area, Costa Rica’, Revista de Biología Tropical, 57(4). Available at: 10.15517/rbt.v57i4.5459.

Calderón-Arguedas, O., Avendaño, A., López-Sánchez, W., and Troyo, A. (2010) ‘Expansion of *Aedes albopictus* skull in Costa Rica’, Rev. Ibero-Latinoam. Parasitol., 69(2), pp. 220–222.

Calderón-Arguedas, O., Troyo, A., Moreira-Soto, R.D., Marín, R., and Taylor, L. (2015) ‘Dengue viruses in *Aedes albopictus* Skuse from a pineapple plantation in Costa Rica’, Journal of Vector Ecology, 40(1), pp. 184–186. Available at: 10.1111/jvec.12149.

Callahan, B.J., McMurdie, P.J., Rosen, M.J., Han, A.W., Johnson, A.J.A., and Holmes, S.P.(2016) ‘DADA2: High-resolution sample inference from Illumina amplicon data’, Nature Methods, 13(7), pp. 581–583. Available at: 10.1038/nmeth.3869.

Câmara, D.C.P., Pinel, C.S., Rocha, G.P., Codeço, C.T., and Honório, N.A. (2020) ‘Diversity of mosquito (Diptera: Culicidae) vectors in a heterogeneous landscape endemic for arboviruses’, Acta Tropica, 212, p. 105715. Available at: 10.1016/j.actatropica.2020.105715.

Canelas, T., Thomsen, E., KAmgang, B., and Kelly-Hope, L.A. (2023) ‘Demographic and environmental factors associated with the distribution of *Aedes albopictus* in Cameroon’, Medical and Veterinary Entomology, 37(1), pp. 143–151. Available at: 10.1111/mve.12619.

Chaverri, L.G., Dillenbeck, C., Lewis, D., Rivera, C., Romero, L.M., and Chaves, L.F. (2018) ‘Mosquito species (Diptera: Culicidae) diversity from ovitraps in a Mesoamerican tropical rainforest’, Journal of Medical Entomology, 55(3), pp. 646–653. Available at: 10.1093/jme/tjx254.

Chaves, A., Piche-Ovares, M., Ibarra-Cerdeña, C.N., Corrales-Aguilar, E., Suzán, G., Moreira-Soto, A., and Gutiérrez-Espeleta, G. (2021) ‘Serosurvey of nonhuman primates in Costa Rica at the human–wildlife interface reveals high exposure to Flaviviruses’, Insects, 12(6), p. 554. Available at: 10.3390/insects12060554.

Chaves, L.S.M., Bergo, E.S., Conn, J.E., Laporta, G.Z., Prist, P.R., and Sallum, M.A.M. (2021) ‘Anthropogenic landscape decreases mosquito biodiversity and drives malaria vector proliferation in the Amazon rainforest’, PLOS ONE, 16(1), p. e0245087. Available at: 10.1371/journal.pone.0245087.

Chazdon, R.L., Harvey, C.A., Komar, O., Griffith, D.M., Ferguson, B.G., Martínez-Ramos, M., *et. al.* (2009) ‘Beyond reserves: A research agenda for conserving biodiversity in human-modified tropical landscapes’, Biotropica, 41(2), pp. 142–153. Available at: 10.1111/j.1744-7429.2008.00471.x.

Coddington, J.A., Agnarrson, I., Miller, J.A., Kuntner, M. and Hormiga, G. (2009) ‘Undersampling Bias: The Null Hypothesis for Singleton Species in Tropical Arthropod Surveys’, Journal of Animal Ecology, 78(3), pp. 573–584.

Coetzee, B.W.T., Gaston, K.J. and Chown, S.L. (2014) ‘Local scale comparisons of biodiversity as a test for global protected area ecological performance: A meta-analysis’, PLOS ONE, 9(8), p. e105824. Available at: 10.1371/journal.pone.0105824.

Daily, G.C., Ehrlich, P.R. and Sánchez-Azofeifa, G.A. (2001) ‘Countryside Biogeography: Use of Human-Dominated Habitats by the Avifauna of Southern Costa Rica’, Ecological Applications, 11(1), pp. 1–13. Available at: 10.1890/1051-0761(2001)011[0001:CBUOHD]2.0.CO;2.

Daugherty, M.P., Alto, B.W. and Juliano, S.A. (2000) ‘Invertebrate carcasses as a resource for competing *Aedes albopictus* and *Aedes aegypti* (Diptera: Culicidae)’, Journal of Medical Entomology, 37(3), pp. 364–372. Available at: 10.1093/jmedent/37.3.364.

Echeverri, A., Smith, J.R., MacArthur-Waltz, D., Lauck, K.S., Anderson, C.B., Monge Vargas, R., *et. al.* (2022) ‘Biodiversity and infrastructure interact to drive tourism to and within Costa Rica’, Proceedings of the National Academy of Sciences, 119(11), p. e2107662119. Available at: 10.1073/pnas.2107662119.

Ferraguti, M., Martínez-de la Puente, J., Roiz, D., Ruiz, S., Soriguer, R., and Figuerola, J. (2016) ‘Effects of landscape anthropization on mosquito community composition and abundance’, Scientific Reports, 6(1), p. 29002. Available at: 10.1038/srep29002.

Fick, S.E. and Hijmans, R.J. (2017) ‘WorldClim 2: New 1-km spatial resolution climate surfaces for global land areas’, International journal of climatology, 37(12), pp. 4302–4315.

Fitzpatrick, M., Mokany, K., Manion, G., Nieto-Lugilde, D., and Ferrier, S. (2022) ‘gdm: Generalized Dissimilarity Modeling’, R package. Available at: https://CRAN.R-project.org/package=gdm.

Frank, H.K., Frishkoff, L.O., Mendenhall, C.D., Daily, G.C., and Hadly, E.A. (2017) ‘Phylogeny, traits, and biodiversity of a Neotropical bat assemblage: Close relatives show similar responses to local deforestation’, The American Naturalist, 190(2), pp. 200–212. Available at: 10.1086/692534.

Franklinos, L.H.V., Jones, K.E., Redding, D.W., and Abubakar, I. (2019) ‘The effect of global change on mosquito-borne disease’, The Lancet Infectious Diseases, 19(9), pp. e302–e312. Available at: 10.1016/S1473-3099(19)30161-6.

Frishkoff, L.O., Ke, A., Martins, I.S., Olimpi, E.M., and Karp, D.S. (2019) ‘Countryside biogeography: The controls of species distributions in human-dominated landscapes’, Current Landscape Ecology Reports, 4(2), pp. 15–30. Available at: 10.1007/s40823-019-00037-5.

Garmendia, A.E., Van Kruiningen, H.J. and French, R.A. (2001) ‘The West Nile virus: Its recent emergence in North America’, Microbes and Infection, 3(3), pp. 223–229. Available at: 10.1016/S1286-4579(01)01374-0.

Giam, X. (2017) ‘Global biodiversity loss from tropical deforestation’, Proceedings of the National Academy of Sciences, 114(23), pp. 5775–5777. Available at: 10.1073/pnas.1706264114.

Gilotra, S.K., Rozeboom, L.E. and Bhattacharya, N.C. (1967) ‘Observations on possible competitive displacement between populations of *Aedes aegypti* Linnaeus and *Aedes albopictus* Skuse in Calcutta’, Bulletin of the World Health Organization, 37(3), pp. 437–446.

Giordano, B.V., Bartlett, S.K., Falcaon, D.A., Lucas, R.P., Tressler, M.J. and Campbell, L.P. (2020) ‘Mosquito Community Composition, Seasonal Distributions, and Trap Bias in Northeastern Florida’, Journal of Medical Entomology, 57(5), pp. 1501–1509. Available at: 10.1093/jme/tjaa053.

Gutman, G., Janetos A. C., Justice, C.O., Moran, E.F., Mustard, J.F., Rindfuss, R.R. et. al. (eds) (2004) Land Change Science: Observing, Monitoring and Understanding Trajectories of Change on the Earth’s Surface. Dordrecht: Springer Netherlands (Remote Sensing and Digital Image Processing). Available at: 10.1007/978-1-4020-2562-4.

Hardulak, L.A., Morinière, J., Hausmann, A., Hendrich, L., Schmidt, S., Doczkal, D., *et. al.* (2020) ‘DNA metabarcoding for biodiversity monitoring in a national park: Screening for invasive and pest species’, Molecular Ecology Resources, 20(6), pp. 1542–1557. Available at: 10.1111/1755-0998.13212.

Heard, S.B. (1994) ‘Pitcher-plant midges and mosquitoes: A processing chain commensalism’, Ecology, 75(6), pp. 1647–1660. Available at: 10.2307/1939625.

Hendershot, J.N., Smith, J.R., Anderson, C.B., Letten, A.D., Frishkoff, L.O., Zook, J.R., et al. (2020) ‘Intensive farming drives long-term shifts in avian community composition’, Nature, 579(7799), pp. 393–396. Available at: 10.1038/s41586-020-2090-6.

Hendy, A., Hernandez-Acosta, E., Valério, D., Fé, N.F., Mendonça, C.R., Costa, E.R., et al. (2023) ‘Where boundaries become bridges: Mosquito community composition, key vectors, and environmental associations at forest edges in the central Brazilian Amazon’, PLOS Neglected Tropical Diseases, 17(4), p. e0011296. Available at: 10.1371/journal.pntd.0011296.

Hijmans, R.J., Van Etten, J., Cheng, J., Mattiuzzi, M., Sumner, M., Greenberg, J.A., Lamigueiro, O.P., Bevan, A., Racine, E.B., Shortridge, A. and Hijmans, M.R.J., 2015. Package ‘raster’. R package, 734, p.473.

Hooke, R., Le B., Martín-Duque, J.F. and Pedraza, J. (2012) ‘Land transformation by humans: A review’, GSA Today, 12(12), pp. 4–10. Available at: 10.1130/GSAT151A.1.

Hoshi, T., Imanishi, N., Higa, Y. and Chaves, L.F. (2014) ‘Mosquito Biodiversity Patterns Around Urban Environments in South-Central Okinawa Island, Japan’, Journal of the American Mosquito Control Association, 30(4), pp. 260–267. Available at: 10.2987/14-6432R.1.

Johnson, M.F., Gómez, A. and Pinedo-Vasquez, M. (2008) ‘Land use and mosquito diversity in the Peruvian Amazon’, Journal of Medical Entomology, 45(6), pp. 1023–1030. Available at: 10.1603/0022-2585(2008)45[1023:LUAMDI]2.0.CO;2.

Jose, S. (2009) ‘Agroforestry for ecosystem services and environmental benefits: An overview’, Agroforestry Systems, 76(1), pp. 1–10. Available at: 10.1007/s10457-009-9229-7.

Karp, D.S., Mendenhall, C.D., Sandí, R.F., Chaumont, N., Ehrlich, P.R., Hadly, E.A., and Daily, G.C. (2013) ‘Forest bolsters bird abundance, pest control and coffee yield’, Ecology Letters, 16(11), pp. 1339–1347. Available at: 10.1111/ele.12173.

Kindt, R. and Coe, R. (2005). Tree diversity analysis. A manual and software for common statistical methods for ecological and biodiversity studies. World Agroforestry Centre (ICRAF). ISBN 92-9059-179-X, http://www.worldagroforestry.org/output/tree-diversity-analysis.

Kirmse, S. (2024) ‘Structure and composition of a canopy-beetle community (Coleoptera) in a Neotropical lowland rainforest in southern Venezuela’, Royal Society Open Science, 11(7), p. 240478. Available at: 10.1098/rsos.240478.

Kraemer, M.U., Sinka, M.E., Duda, K.A., Mylne, A.Q.N., Shearer, F.M., Barker, C.M., *et. al.* (2015) ‘The global distribution of the arbovirus vectors *Aedes aegypti* and *Ae. albopictus*’, eLife, 4, p. e08347. Available at: 10.7554/eLife.08347.

Laird, M. (1988) The natural history of larval mosquito habitats. Academic Press Ltd, London, UK. ISBN: 9780124340053. Available at: http://www.cabdirect.org/cabdirect/abstract/19892056420 (Accessed: 5 July 2021).

Langhans, K.E., Schmitt, R.J.P., Chaplin-Kramer, R., Anderson, C.B., Vargas Bolaños, C., Vargas Cabezas, F., et al. (2022) ‘Modeling multiple ecosystem services and beneficiaries of riparian reforestation in Costa Rica’, Ecosystem Services, 57, p. 101470. Available at: 10.1016/j.ecoser.2022.101470.

LaPointe, D.A., Atkinson, C.T. and Samuel, M.D. (2012) ‘Ecology and conservation biology of avian malaria: Ecology of avian malaria’, Annals of the New York Academy of Sciences, 1249(1), pp. 211–226. Available at: 10.1111/j.1749-6632.2011.06431.x.

Lemon, S.M., Sparling, P.F., Hamburg, M.A., Relman, D.A., Choffnes, E.R., and Mack, A. (eds) (2008) Vector-borne diseases - understanding the environmental, human health, and ecological connections: workshop summary. Washington, D.C.: National Academies Press.

Lichtenberg, E.M., Kennedy, C.M., Kremen, C., Batáry, P., Berendse, F., Bommarco, R., *et. al.* (2017) ‘A global synthesis of the effects of diversified farming systems on arthropod diversity within fields and across agricultural landscapes’, Global Change Biology, 23(11), pp. 4946– 4957. Available at: 10.1111/gcb.13714.

Long, J.A. and Long, M.J.A., 2019. Package ‘interactions’. *See* https://interactions.jacob-long.com.

Maxwell, S.L., Fuller, R.A., Brooks, T.M., and Watson, J.E.M. (2016) ‘Biodiversity: The ravages of guns, nets and bulldozers’, Nature, 536(7615), pp. 143–145. Available at: 10.1038/536143a.

Medlin, S., Deardorff, E.R., Hanley, C.S., Vergneau-Grosset, C., Siudak-Campfield, A., Dallwig, R., *et. al.* (2016) ‘Serosurvey of selected arboviral pathogens in free-ranging, two-toed sloths (*Choloepus hoffmanni*) and three-toed sloths (*Bradypus variegatus*) in Costa Rica, 2005–07’, Journal of wildlife diseases, 52(4), pp. 883–892.

Mendenhall, C.D., Frishkoff, L.O., Santos-Barrera, G., Pacheco, J., Mesfun, E., Quijano, F. M., *et. al.* (2014) ‘Countryside biogeography of Neotropical reptiles and amphibians’, Ecology, 95(4), pp. 856–870. Available at: 10.1890/12-2017.1.

Mendenhall, C.D., Shields-Estrada, A., Krishnaswami, A.J., and Daily, G.C. (2016) ‘Quantifying and sustaining biodiversity in tropical agricultural landscapes’, Proceedings of the National Academy of Sciences, 113(51), pp. 14544–14551. Available at: 10.1073/pnas.1604981113.

Meyer Steiger, D.B., Ritchie, S.A. and Laurance, S.G.W. (2016) ‘Mosquito communities and disease risk influenced by land use change and seasonality in the Australian tropics’, Parasites & Vectors, 9(1), p. 387. Available at: 10.1186/s13071-016-1675-2.

Millennium ecosystem assessment, M. (2005) Ecosystems and human well-being. Island press Washington, DC.

Mordecai, E.A., Caldwell, J.M., Grossman, M.K., Lippi, C.A., Johnson, L.R., Neira, M., et al. (2019) ‘Thermal biology of mosquito-borne disease’, Ecology Letters. Edited by J. (Jeb) Byers, p. ele.13335. Available at: 10.1111/ele.13335.

Muturi, E.J., Shililu, J., Jacob, B., Gu, W., Githure, J., and Novak, R. (2006) ‘Mosquito species diversity and abundance in relation to land use in a riceland agroecosystem in Mwea, Kenya’, Journal of Vector Ecology, 31(1), pp. 129–137. Available at: 10.3376/1081-1710(2006)31[129:MSDAAI]2.0.CO;2.

Niebylski, M.L., Savage, H.M., Nasci, R.S., and Craig, G.B. (1994) ‘Blood hosts of *Aedes albopictus* in the United States’, Journal of the American Mosquito Control Association, 10(3), pp. 447–450.

Norris, K. (2008) ‘Agriculture and biodiversity conservation: opportunity knocks’, Conservation Letters, 1(1), pp. 2–11. Available at: 10.1111/j.1755-263X.2008.00007.x.

Novotný, V. and Basset, Y. (2000) ‘Rare species in communities of tropical insect herbivores: pondering the mystery of singletons’, Oikos, 89(3), pp. 564–572. Available at: 10.1034/j.1600-0706.2000.890316.x.

Oksanen, J., Blanchet, F.G., Kindt, R., Legendr, P., Minchin, P.R., O’Hara, R.B., et al. (2013) ‘Package “vegan”’, Community ecology package, version, 2(9), pp. 1–295.

Pagès, H., Aboyoun, P., Gentleman, R., and DebRoy, S. (2022) ‘Biostrings: Efficient manipulation of biological strings’. (R package). Available at: https://bioconductor.org/packages/Biostrings.

Pereira, H.M., Leadley, P.W., Proença, V., Alkemade, R., Scharlemann, J.P.W., Fernandez-Manjarrés, J.F., et al. (2010) ‘Scenarios for global biodiversity in the 21st Century’, Science, 330(6010), pp. 1496–1501. Available at: 10.1126/science.1196624.

Pereira dos Santos, T., Roiz, D., Santos de Abreau, F.V., Luz, S.L.B., Santalucia, M., Jiolle, D., et al. (2018) ‘Potential of Aedes albopictus as a bridge vector for enzootic pathogens at the urban-forest interface in Brazil’, Emerging Microbes & Infections, 7(1), pp. 1–8. Available at: 10.1038/s41426-018-0194-y.

Perrin, A., Glaizot, O. and Christe, P. (2022) ‘Worldwide impacts of landscape anthropization on mosquito abundance and diversity: A meta-analysis’, Global Change Biology, 28(23), pp. 6857– 6871. Available at: 10.1111/gcb.16406.

Piaggio, M., Guzman, M., Pacay, E., Robalino, J. and Ricketts, T. (2024) ‘Forest Cover and Dengue in Costa Rica: Panel Data Analysis of the Effects of Forest Cover Change on Hospital Admissions and Outbreaks’, Environmental and Resource Economics, 87(8), pp. 2095–2114. Available at: 10.1007/s10640-024-00853-2.

Piche-Ovares, M., Romero-Vega, M., Vargas-González, D., Murillo, D.F.P, Soto-Garita, C., Francisco-Llamas, J., et. al. (2023) ‘Serosurvey in two dengue hyperendemic areas of Costa Rica evidence active circulation of WNV and SLEV in peri-domestic and domestic animals and in humans’, Pathogens, 12(1), p. 7. Available at: 10.3390/pathogens12010007.

Poulin, B., Lefebvre, G. and Paz, L. (2010) ‘Red flag for green spray: adverse trophic effects of Bti on breeding birds’, Journal of Applied Ecology, 47(4), pp. 884–889. Available at: 10.1111/j.1365-2664.2010.01821.x.

Prevedello, J.A., Winck, G.R., Weber, M.M., Nichols, E., Sinervo, B., and Xu, Y. (2019) ‘Impacts of forestation and deforestation on local temperature across the globe’, PLOS ONE. Edited by Y. Xu, 14(3), p. e0213368. Available at: 10.1371/journal.pone.0213368.

Ratnasingham, S. and Hebert, P.D.N. (2007) ‘bold: The Barcode of Life data system (http://www.barcodinglife.org)’, Molecular Ecology Notes, 7(3), pp. 355–364. Available at: 10.1111/j.1471-8286.2007.01678.x.

Reisen, W.K. (2003) ‘Epidemiology of St. Louis encephalitis virus’, Advances in virus research, 61, pp. 139–184.

Responte, M. and Nuneza, O. (2016) ‘Diversity of pholcid spiders (Araneae:Pholcidae) in selected areas of Southeastern Mindanao, Philippines’, ELBA Bioflux, 8, pp. 7–17.

Rezza, G. (2012) ‘*Aedes albopictus* and the reemergence of dengue’, BMC Public Health, 12(1), p. 72. Available at: 10.1186/1471-2458-12-72.

Rhodes, C.G., Loaiza, J.R., Romero, L.M., Gutiérrez Alvarado, J.M., Delgado, G., Rojas Salas, O., et al. (2022) ‘Anopheles albimanus (Diptera: Culicidae) Ensemble Distribution Modeling: Applications for Malaria Elimination’, Insects, 13(3), p. 221. Available at: 10.3390/insects13030221.

Richards, S.L., Ponnusamy, L., Unnasch, T.R., Hassan, H.K., and Apperson, C.S. (2006) ‘Host-feeding patterns of *Aedes albopictus* (Diptera: Culicidae) in relation to availability of human and domestic animals in suburban landscapes of central North Carolina’, Journal of Medical Entomology, 43(3), pp. 543–551. Available at: 10.1093/jmedent/43.3.543.

Ricketts, T.H., Daily, G.C., Ehrlich, P.R., and Michener, C.D. (2004) ‘Economic value of tropical forest to coffee production’, Proceedings of the National Academy of Sciences, 101(34), pp. 12579–12582. Available at: 10.1073/pnas.0405147101.

Ripley, B., Venables, B., Bates, D.M., Hornik, K., Gebhardt, A., Firth, D. and Ripley, M.B., 2013. Package ‘mass’. Cran r, 538, pp.113–120.

Rojas-Araya, D., Marín-Rodríguez, R., Gutiérrez-Alvarado, M., Romero-Vega, L.M., Calderón-Arguedas, O., and Troyo, A. (2017) ‘New records of *Aedes albopictus* (Skuse) in four locations of Costa Rica’, Revista Biomédica, 28(2), pp. 79–88. Available at: 10.32776/revbiomed.v28i2.575.

Romero-Vega, L.M., Piche-Ovares, M., Soto-Garita, C., Barantes Murillo, D.F., Chaverri, L.G., Alfaro-Alarcón, A., et al. (2023) ‘Seasonal changes in the diversity, host preferences and infectivity of mosquitoes in two arbovirus-endemic regions of Costa Rica’, Parasites & Vectors, 16(1), p. 34. Available at: 10.1186/s13071-022-05579-y.

Sala, O.E. (2000) ‘Global biodiversity scenarios for the year 2100’, Science, 287(5459), pp. 1770–1774. Available at: 10.1126/science.287.5459.1770.

Sánchez-Azofeifa, G.A., Pfaff, A., Robalino, J.A., and Boomhower, J.P. (2007) ‘Costa Rica’s Payment for Environmental Services program: intention, implementation, and impact’, Conservation Biology, 21(5), pp. 1165–1173. Available at: 10.1111/j.1523-1739.2007.00751.x.

da Silva Pessoa Vieira, C.J., Steiner São Bernardo, C., Ferriera da Silva, D.J., Rigotti Kubiszeski, J., Serpa Barreto, E., de Oliveira Monteiro, H.A., et al. (2022) ‘Land-use effects on mosquito biodiversity and potential arbovirus emergence in the Southern Amazon, Brazil’, Transboundary and Emerging Diseases, 69(4), pp. 1770–1781. Available at: 10.1111/tbed.14154.

Simmons, C.P., Farrar, J.J., van Vinh Chau, N., and Wills, B. (2012) ‘Dengue’, New England Journal of Medicine, 366(15), pp. 1423–1432. Available at: 10.1056/NEJMra1110265.

Smith, R.J. (2022) R package ‘ecole’, phytomosaic/ecole: ecole: School of Ecology Package. Available at: https://rdrr.io/github/phytomosaic/ecole/.

Smith, T.J. and Mayfield, M.M. (2015) ‘Diptera species and functional diversity across tropical Australian countryside landscapes’, Biological Conservation, 191, pp. 436–443. Available at: 10.1016/j.biocon.2015.07.035.

Sokolow, S.H., Nova, N., Jones, I.J., Wood, C.L., Lafferty, K.D., Garchitorena, A., et al. (2022) ‘Ecological and socioeconomic factors associated with the human burden of environmentally mediated pathogens: a global analysis’, The Lancet Planetary Health, 6(11), pp. e870–e879. Available at: 10.1016/S2542-5196(22)00248-0.

Thongsripong, P., Green, A., Kittayapong, P., Kapan, D., Wilcox, B., and Bennett, S. (2013) ‘Mosquito vector diversity across habitats in central Thailand endemic for dengue and other arthropod-borne diseases’, PLOS Neglected Tropical Diseases, 7(10), p. e2507. Available at: 10.1371/journal.pntd.0002507.

Thormann, B., Ahrens, D., Armijos, D.M., Peters, M.K., Wagner, T. and Wägele, J.W. (2016) ‘Exploring the Leaf Beetle Fauna (Coleoptera: Chrysomelidae) of an Ecuadorian Mountain Forest Using DNA Barcoding’, PLOS ONE, 11(2), p. e0148268. Available at: 10.1371/journal.pone.0148268.

Troyo, A., Porcelain, S.L., Calderón-Arguedas, O., Chadee, D.D., and Beier, J.C. (2006) ‘Dengue in Costa Rica: the gap in local scientific research’, Revista Panamericana de Salud Pública, 20, pp. 350–360. Available at: 10.1590/S1020-49892006001000012.

Troyo, A., Calderón-Arguedas, O., Fuller, D.O., Solano, M.E., Avendaño, A., et al. (2008) ‘Seasonal profiles of *Aedes aegypti* (Diptera: Culicidae) larval habitats in an urban area of Costa Rica with a history of mosquito control’, Journal of Vector Ecology, 33(1), pp. 76–88. Available at: 10.3376/1081-1710(2008)33[76:SPOAAD]2.0.CO;2.

Troyo, A., Fuller, D.O., Calderón-Arguedas, O., Solano, M.E., and Beier, J.C. (2009) ‘Urban structure and dengue incidence in Puntarenas, Costa Rica’, Singapore Journal of Tropical Geography, 30(2), pp. 265–282. Available at: 10.1111/j.1467-9493.2009.00367.x.

Tsuda, Y., Suwonkerd, W., Chawprom, S., Prajakwong, S., and Takagi, M. (2006) ‘Different spatial distribution of *Aedes aegypti* and *Aedes albopictus* along an urban-rural gradient and the relating environmental factors examined in three villages in northern Thailand’, Journal of the American Mosquito Control Association, 22(2), pp. 222–228. Available at: 10.2987/8756-971X(2006)22[222:DSDOAA]2.0.CO;2.

Wickham, H (2007). “Reshaping data with the reshape package.” Journal of Statistical Software, 21(12), 1–20. http://www.jstatsoft.org/v21/i12/.

Wickham, H., Averick, M., Bryan, J., Chang, W., Mcgowan, L.D., François, R., et al. (2019) ‘Welcome to the Tidyverse’, Journal of Open Source Software, 4(43), p. 1686. Available at: 10.21105/joss.01686.

Wickham H, François R, Henry L, Müller K, Vaughan D (2023). dplyr: A Grammar of Data Manipulation. R package version 1.1.4, https://github.com/tidyverse/dplyr, https://dplyr.tidyverse.org.

Wilke, C.O., Wickham, H. and Wilke, M.C.O., 2019. Package ‘cowplot’. Streamlined plot theme and plot annotations for ‘ggplot2, 1.

World Health Organization (1994) ‘Outbreak of classic dengue, Costa Rica’, Weekly Epidemiological Record, 69, pp. 85–86.

World Health Organization (2021) Dengue data application. Available at: https://ntdhq.shinyapps.io/dengue5.

Wright, E. (2016) ‘Using DECIPHER v2.0 to analyze big biological sequence data in R’, The R Journal, 8(1), pp. 352–359.

Young, K.I., Mundis, S., Widen, S.G., Wood, T.G., Tesh, R.B., Cardosa, J., et al. (2017) ‘Abundance and distribution of sylvatic dengue virus vectors in three different land cover types in Sarawak, Malaysian Borneo’, Parasites & Vectors, 10(1), p. 406. Available at: 10.1186/s13071-017-2341-z.

Zimmerman, R.H. (1992) ‘Ecology of malaria vectors in the Americas and future direction’, Memórias do Instituto Oswaldo Cruz, 87, pp. 371–383. Available at: 10.1590/S0074-02761992000700064

