## Supplemental Information for "Local tree cover predicts mosquito species richness and disease vector presence in a tropical countryside landscape"

**Supplemental figures and tables for "Local tree cover predicts mosquito species richness and disease vector presence in a tropical countryside landscape"**
August 31, 2024

**Figure S1. Species accumulation curves for each land use type were not asymptotic.** Gray shading shows 95% confidence intervals.

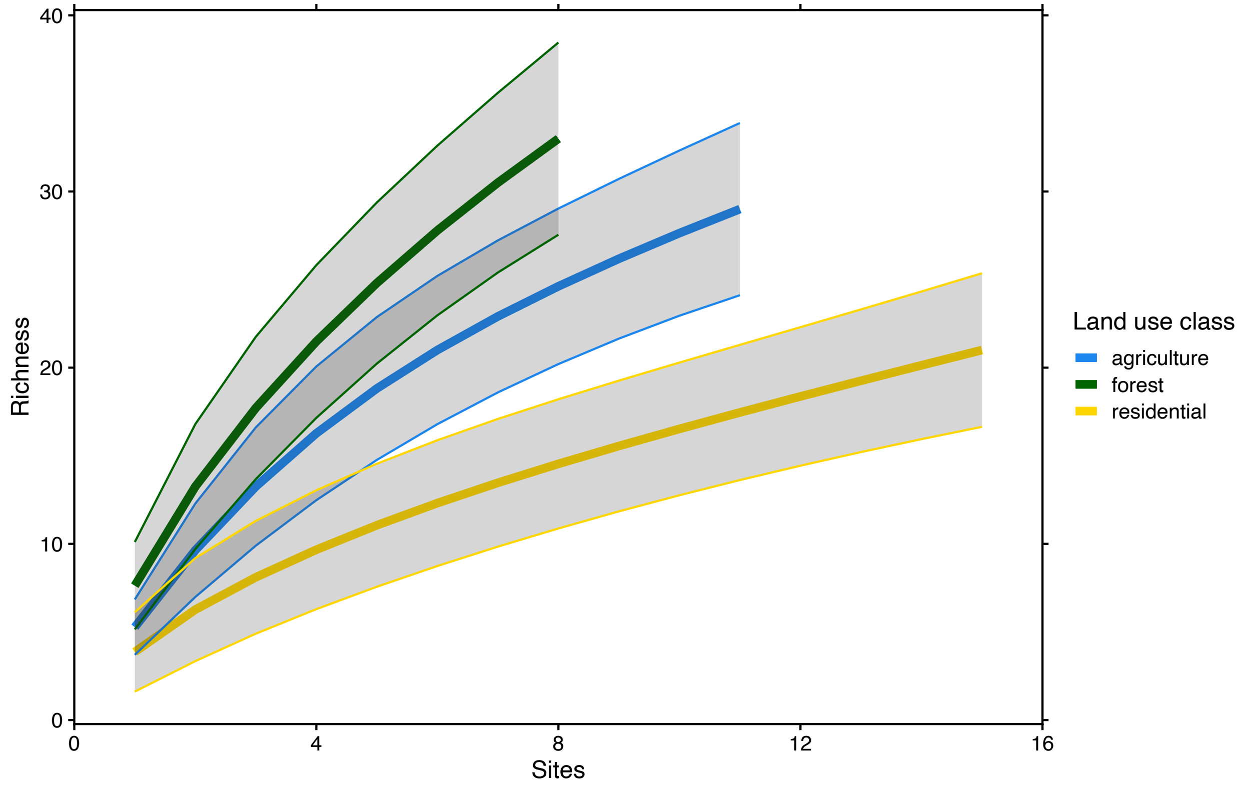

**Figure S2. At the 1000 m spatial scale, species richness increased with percent tree cover only where mean annual temperatures were low to intermediate. In the warmest climates surveyed, species richness was low even at high tree cover.** Points representing each survey site are colored by mean annual temperature. The blue regression line shows the results of a negative binomial GLM modeling the effect of tree cover on species richness in cooler climates (mean annual temperature ≤ 24 ºC; estimate = $2.25 \times 10^{-2}$, SE = $6.92 \times 10^{-3}$, Z = 3.26, p = $1.13 \times 10^{-3}$), and the red line shows the results for warmer climates (mean annual temperature > 24 ºC; -estimate = $-3.7 \times 10^{-2}$, SE = $1.46 \times 10^{-2}$, Z = -2.53, p = $1.13 \times 10^{-2}$).

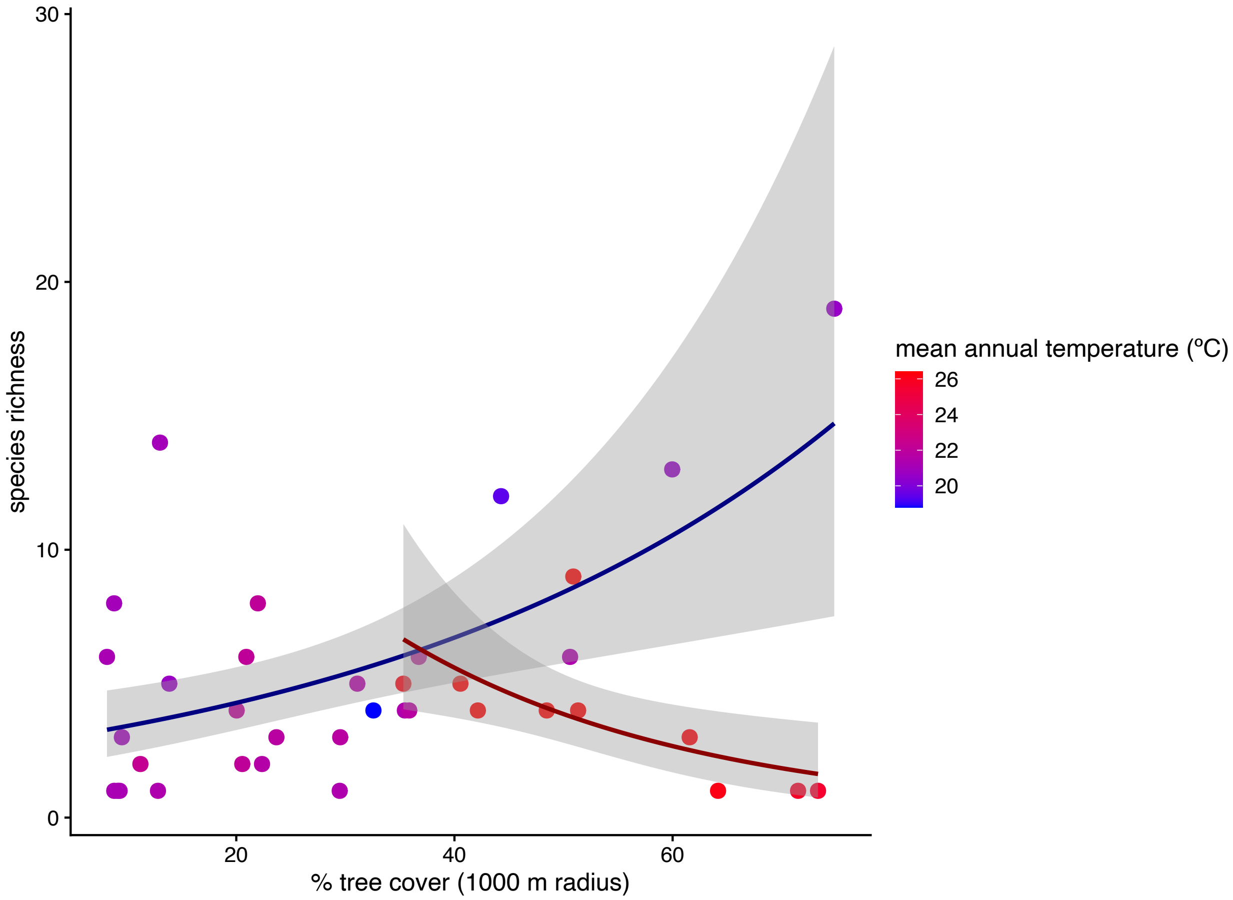

**Figure S3. At the 1000 m spatial scale, *Ae. albopictus* occurrence is negatively correlated with tree cover, and positively correlated with mean annual temperature.**

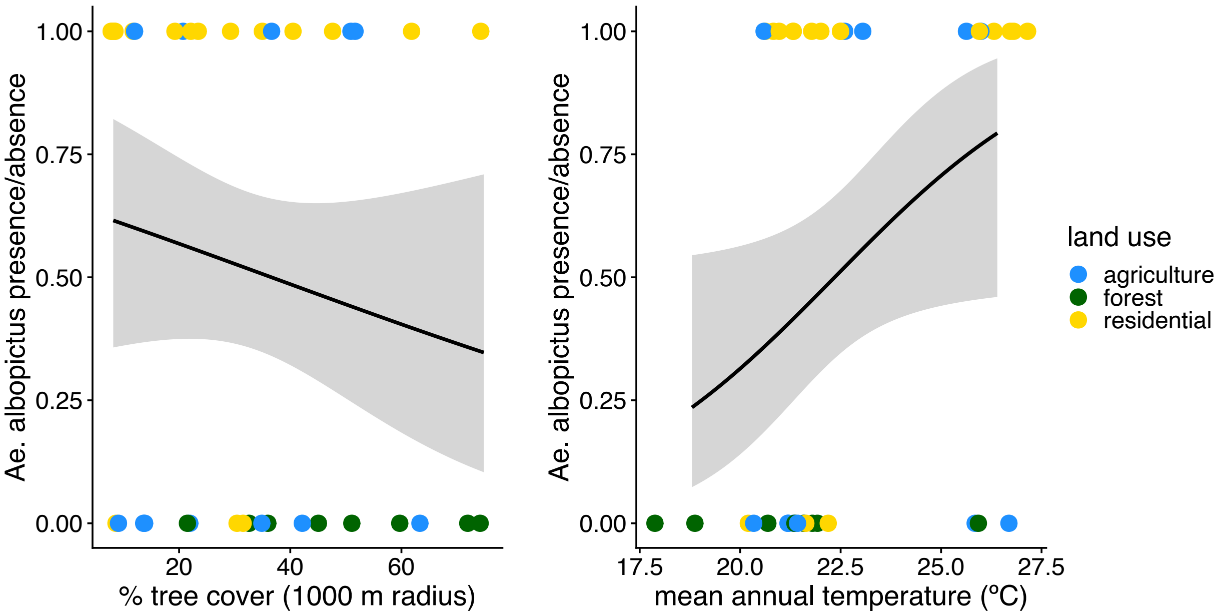

**Figure S4. NMDS ordination plot of community similarity with points labeled by land use subtype.** Red boxes indicate the four sites that were sampled only once, all of which were located in the geographically distinct Pavones district.

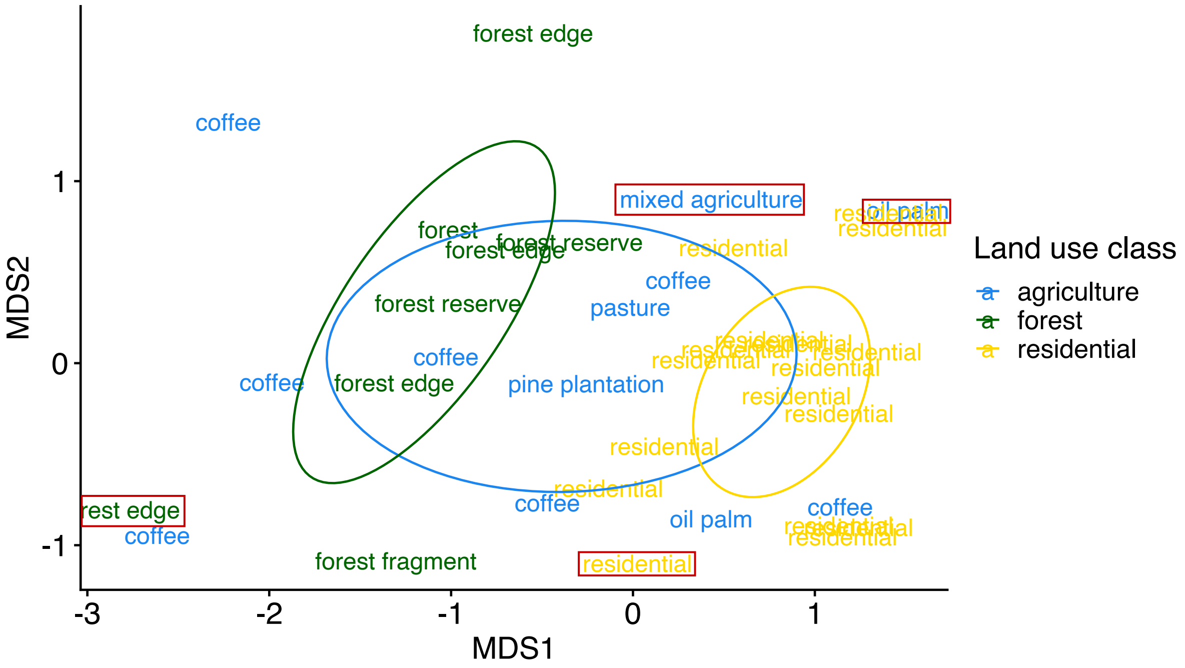

**Table S1. Field survey site collection details, grouped by canton, district, and land use type.**

| Canton | District | Land use | Site | Collection dates (2017) | Total mosquitoes sampled | Morphologically identified *Aedes albopictus* |
|---|---|---|---|---|---|---|
| **Corredores** | *Ciudad Neily* | agriculture | OP03B | 6/30, 7/20 | 32 | 16 |
| | | | OP03 | 6/30, 7/20 | 29 | 12 |
| | | residential | CN04 | 6/30, 7/20 | 210 | 1 |
| | | | CN06 | 6/30, 7/20 | 12 | 5 |
| | | | OP01 | 6/30, 7/20 | 11 | 1 |
| | | | CN03 | 6/30, 7/20 | 5 | 3 |
| | | | | | *Total = 299* | *Total = 38* |
| **Coto Brus** | *Copabuena* | agriculture | SAFR | 7/3, 7/11 | 7 | 0 |
| | | | ELPU | 6/26, 7/11 | 5 | 0 |
| | | residential | CB03 | 6/28, 7/10 | 37 | 16 |
| | | | CB04 | 7/28, 8/3 | 20 | 2 |
| | | | CB05 | 7/10, 8/3 | 5 | 2 |
| | | | CB01 | 6/28, 7/10 | 4 | 2 |
| | | | CB02 | 6/28, 7/10 | 4 | 0 |
| | | | | | *Total = 82* | *Total = 22* |
| | *Sabalito* | agriculture | LOAN | 6/26, 8/1 | 204 | 0 |
| | | | VALE | 6/27, 7/24 | 7 | 0 |
| | | | FELO | 6/27, 7/24 | 4 | 2 |
| | | | SATE | 6/22, 7/13 | 4 | 2 |
| | | forest | QUST | 6/22, 7/13 | 3 | 0 |
| | | residential | SL02 | 7/22, 7/7 | 26 | 14 |
| | | | SL01 | 7/22, 7/7 | 20 | 4 |
| | | | | | *Total = 268* | *Total = 22* |
| | *San Vito* | agriculture | PINO | 6/21, 7/13 | 14 | 6 |
| | | | ISLA | 6/21, 7/13 | 2 | 0 |
| | | forest | MELI | 6/25, 7/31 | 244 | 0 |

| | | | | | | |
|---|---|---|---|---|---|---|
| | | | RIJA | 6/25, 7/31 | 172 | 0 |
| | | | GAPA | 6/23, 7/18 | 97 | 0 |
| | | | FRAG5 | 6/20, 7/18 | 20 | 0 |
| | | | FILA | 6/20, 7/17 | 16 | 0 |
| | | | GABO | 6/23, 7/18 | 12 | 0 |
| | | residential | SV04 | 7/3, 7/25 | 12 | 6 |
| | | | SV06 | 6/20, 7/25 | 12 | 0 |
| | | | SV02 | 6/21, 7/3 | 8 | 1 |
| | | | SV05 | 6/21, 7/3 | 7 | 1 |
| | | | SV01 | 6/23, 7/3 | 3 | 0 |
| | | | | | *Total = 619* | *Total = 14* |
| **Golfito** | *Pavones* | agriculture | PV02 | 7/5 | 2 | 0 |
| | | | PV04 | 7/5 | 2 | 0 |
| | | forest | PV05 | 7/5 | 1 | 0 |
| | | residential | PV01 | 7/5 | 11 | 3 |
| | | | | | *Total = 16* | *Total = 3* |

**Table S2. Details of mosquito taxa present at each survey site, grouped by canton and land use type.**

| Canton | Land use | Site | Trap nights | OTU | Species |
|---|---|---|---|---|---|
| Coto Brus | residential | CB01 | 2 | AA | *Aedes albopictus* |
| | | CB02 | 2 | OTU23 | *Culex quinquefasciatus* |
| | | CB03 | 2 | OTU04 | *Aedes albopictus* |
| | | | | AA | *Aedes albopictus* |
| | | CB04 | 2 | OTU03 | *Culex* sp. 1 |
| | | | | OTU04 | *Aedes albopictus* |
| | | | | OTU07 | *Limatus durhamii* |
| | | | | OTU09 | *Culex nigripalpus* |
| | | | | OTU11 | *Aedes albopictus* |
| | | | | OTU22 | *Culex lactator* |
| | | | | OTU23 | *Culex quinquefasciatus* |
| | | | | AA | *Aedes albopictus* |
| | | CB05 | 2 | OTU04 | *Aedes albopictus* |
| | | | 2 | OTU05 | *Wyeomyia adelpha/guatemala* |
| | | | 2 | OTU07 | *Limatus durhamii* |
| | | | 2 | OTU08 | *Orthopodomyia sp.* |
| | | | 2 | OTU18 | *Orthopodomyia sp.* |
| | | | 2 | OTU19 | *Orthopodomyia sp.* |
| | | | 2 | OTU20 | *Culex coronator* |
| | | | 2 | OTU22 | *Culex lactator* |
| | | | 2 | OTU30 | *Limatus asulleptus* |
| | | | 2 | OTU51 | *Orthopodomyia sp.* |
| | | | 2 | OTU55 | *Orthopodomyia MBI-02* |
| | | | 2 | AA | *Aedes albopictus* |
| | | SL01 | 2 | OTU04 | *Aedes albopictus* |
| | | | | OTU11 | *Aedes albopictus* |
| | | | | OTU23 | *Culex quinquefasciatus* |
| | | | | OTU26 | *Culex interrogator* |
| | | | | OTU34 | *Culex mollis* |
| | | | | AA | *Aedes albopictus* |
| | | SL02 | 2 | OTU23 | *Culex quinquefasciatus* |
| | | | | AA | *Aedes albopictus* |
| | | SV01 | 2 | OTU23 | *Culex quinquefasciatus* |
| | | SV02 | 2 | OTU04 | *Aedes albopictus* |
| | | | | OTU22 | *Culex lactator* |
| | | | | OTU23 | *Culex quinquefasciatus* |

| | | | | |
|---|---|---|---|---|
| | | | OTU57 | *Culex quinquefasciatus* |
| | | | AA | *Aedes albopictus* |
| | SV04 | 2 | OTU05 | *Wyeomyia adelpha/guatemala* |
| | | | OTU07 | *Limatus durhamii* |
| | | | OTU23 | *Culex quinquefasciatus* |
| | | | OTU30 | *Limatus asulleptus* |
| | | | OTU33 | *Wyeomyia aporonoma* |
| | | | OTU40 | *Wyeomyia undulata* |
| | | | OTU72 | *Coquillettidia nigricans* |
| | | | AA | *Aedes albopictus* |
| | SV05 | 2 | OTU12 | *Wyeomyia complosa* |
| | | | OTU23 | *Culex quinquefasciatus* |
| | | | AA | *Aedes albopictus* |
| | SV06 | 2 | OTU07 | *Limatus durhamii* |
| | | | OTU09 | *Culex nigripalpus* |
| | | | OTU22 | *Culex lactator* |
| | | | OTU23 | *Culex quinquefasciatus* |
| | | | OTU72 | *Coquillettidia nigricans* |
| agriculture | ELPU | 2 | OTU03 | *Culex* sp. 1 |
| | | | OTU15 | *Trichoprosopon digitatum* |
| | | | OTU20 | *Culex coronator* |
| | | | OTU35 | *Trichoprosopon pallidiventer* |
| | | | OTU65 | unidentified Culicidae 4 |
| | ISLA | 2 | OTU15 | *Trichoprosopon digitatum* |
| | | | OTU17 | *Haemagogus* sp. |
| | | | OTU27 | *Haemagogus* sp. |
| | LOAN | 2 | OTU03 | *Culex* sp. 1 |
| | | | OTU05 | *Wyeomyia adelpha/guatemala* |
| | | | OTU06 | *Culex nigripalpus* |
| | | | OTU07 | *Limatus durhamii* |
| | | | OTU09 | *Culex nigripalpus* |
| | | | OTU12 | *Wyeomyia complosa* |
| | | | OTU13 | *Aedes allotecnon* |
| | | | OTU14 | *Wyeomyia sp. RCH-1* |
| | | | OTU15 | *Trichoprosopon digitatum* |
| | | | OTU21 | *Psorophora ferox* |
| | | | OTU30 | *Limatus asulleptus* |
| | | | OTU41 | *Aedes* sp. 1 |

| | | | | |
|---|---|---|---|---|
| | | | OTU45 | *Psorophora funiculus* |
| | | | OTU52 | *Wyeomyia* sp. 2 |
| | | | OTU54 | *Aedes allotecnon* |
| | | | OTU64 | unidentified Culicidae 3 |
| | FELO | 2 | AA | *Aedes albopictus* |
| | PINO | 2 | OTU05 | *Wyeomyia adelpha/guatemala* |
| | | | OTU06 | *Culex nigripalpus* |
| | | | OTU12 | *Wyeomyia complosa* |
| | | | OTU30 | *Limatus asulleptus* |
| | | | AA | *Aedes albopictus* |
| | | | AG | *Aedes aegypti* |
| | SAFR | 2 | OTU05 | *Wyeomyia adelpha/guatemala* |
| | | | OTU06 | *Culex nigripalpus* |
| | | | OTU23 | *Culex quinquefasciatus* |
| | SATE | 2 | OTU01 | *Aedes angustivittatus* |
| | | | OTU04 | *Aedes albopictus* |
| | | | OTU07 | *Limatus durhamii* |
| | | | OTU14 | *Wyeomyia sp. RCH-1* |
| | | | OTU15 | *Trichoprosopon digitatum* |
| | | | OTU17 | *Haemagogus* sp. |
| | | | OTU27 | *Haemagogus* sp. |
| | | | AA | *Aedes albopictus* |
| | VALE | 2 | OTU14 | *Wyeomyia sp. RCH-1* |
| | | | OTU21 | *Psorophora ferox* |
| | | | OTU46 | *Haemagogus janthinomys* |
| | | | OTU48 | *Psorophora ferox* |
| | | | OTU62 | *Aedes infirmatus* |
| forest | FILA | 2 | OTU03 | *Culex* sp. 1 |
| | | | OTU05 | *Wyeomyia adelpha/guatemala* |
| | | | OTU06 | *Culex nigripalpus* |
| | | | OTU10 | *Culex restrictor* |
| | | | OTU12 | *Wyeomyia complosa* |
| | | | OTU13 | *Aedes allotecnon* |
| | | | OTU31 | *Aedes trivittatus* |
| | | | OTU41 | *Aedes* sp. 1 |
| | | | OTU42 | *Culex* sp. 2 |
| | | | OTU59 | *Aedes* sp. 1 |
| | | | OTU61 | unidentified Culicidae 2 |

| | | | |
|---|---|---|---|
| | | OTU65 | unidentified Culicidae 4 |
| | | AG | *Aedes aegypti* |
| FRAG5 | 2 | OTU05 | *Wyeomyia adelpha/guatemala* |
| | | OTU06 | *Culex nigripalpus* |
| | | OTU12 | *Wyeomyia complosa* |
| | | OTU46 | *Haemagogus janthinomys* |
| GABO | 2 | OTU05 | *Wyeomyia adelpha/guatemala* |
| | | OTU06 | *Culex nigripalpus* |
| | | OTU15 | *Trichoprosopon digitatum* |
| | | OTU35 | *Trichoprosopon pallidiventer* |
| | | OTU47 | *Trichoprosopon sp. 3 DJS-2020* |
| | | OTU58 | *Trichoprosopon sp.* |
| GAPA | 2 | OTU12 | *Wyeomyia complosa* |
| | | OTU13 | *Aedes allotecnon* |
| | | OTU47 | *Trichoprosopon sp. 3 DJS-2020* |
| | | OTU50 | *Aedes sexlineatus* |
| MELI | 2 | OTU01 | *Aedes angustivittatus* |
| | | OTU03 | *Culex* sp. 1 |
| | | OTU05 | *Wyeomyia adelpha/guatemala* |
| | | OTU06 | *Culex nigripalpus* |
| | | OTU10 | *Culex restrictor* |
| | | OTU12 | *Wyeomyia complosa* |
| | | OTU13 | *Aedes allotecnon* |
| | | OTU20 | *Culex coronator* |
| | | OTU27 | *Haemagogus sp.* |
| | | OTU28 | *Culex erythrothorax* |
| | | OTU29 | *unidentified Culicidae 1* |
| | | OTU30 | *Limatus asulleptus* |
| | | OTU34 | *Culex mollis* |
| | | OTU35 | *Trichoprosopon pallidiventer* |
| | | OTU41 | *Aedes* sp. 1 |
| | | OTU43 | *Aedes angustivittatus* |
| | | OTU48 | *Psorophora ferox* |
| | | OTU51 | *Orthopodomyia* sp. |
| | | OTU66 | *Wyeomyia luteoventralis* |

| | | | | | |
|---|---|---|---|---|---|
| | | | | OTU67 | *Trichoprosopon sp. 5 DJS-2020* |
| | | QUST | 2 | OTU05 | *Wyeomyia adelpha/guatemala* |
| | | | | OTU69 | *Sabethes undosus* |
| | | RIJA | 2 | OTU05 | *Wyeomyia adelpha/guatemala* |
| | | | | OTU06 | *Culex nigripalpus* |
| | | | | OTU09 | *Culex nigripalpus* |
| | | | | OTU12 | *Wyeomyia complosa* |
| | | | | OTU13 | *Aedes allotecnon* |
| | | | | OTU21 | *Psorophora ferox* |
| | | | | OTU23 | *Culex quinquefasciatus* |
| | | | | OTU24 | *Wyeomyia adelpha/guatemala* |
| | | | | OTU30 | *Limatus asulleptus* |
| | | | | OTU35 | *Trichoprosopon pallidiventer* |
| | | | | OTU36 | *Aedes ferox* |
| | | | | OTU41 | *Aedes* sp. 1 |
| | | | | OTU43 | *Aedes angustivittatus* |
| | | | | OTU47 | *Trichoprosopon sp. 3 DJS-2020* |
| | | | | OTU48 | *Psorophora ferox* |
| | | | | OTU63 | *Psorophora albipes* |
| Corredores | residential | CN03 | 2 | OTU23 | *Culex quinquefasciatus* |
| | | | | AA | *Aedes albopictus* |
| | | CN03 | 2 | AG | *Aedes aegypti* |
| | | CN04 | 2 | OTU04 | *Aedes albopictus* |
| | | | | OTU06 | *Culex nigripalpus* |
| | | | | OTU07 | *Limatus durhamii* |
| | | | | OTU11 | *Aedes albopictus* |
| | | | | OTU23 | *Culex quinquefasciatus* |
| | | | | OTU30 | *Limatus asulleptus* |
| | | | | OTU53 | *Aedes albopictus* |
| | | | | AA | *Aedes albopictus* |
| | | CN06 | 2 | AA | *Aedes albopictus* |
| | | OP01 | 2 | OTU06 | *Culex nigripalpus* |
| | | | | OTU22 | *Culex lactator* |
| | | | | OTU23 | *Culex quinquefasciatus* |
| | | | | AA | *Aedes albopictus* |
| | agriculture | OP03 | 2 | OTU01 | *Aedes angustivittatus* |

| | | | | | |
|---|---|---|---|---|---|
| | | | | OTU06 | *Culex nigripalpus* |
| | | | | OTU21 | *Psorophora ferox* |
| | | | | OTU23 | *Culex quinquefasciatus* |
| | | | | OTU25 | *Mansonia titillans* |
| | | | | OTU28 | *Culex erythrothorax* |
| | | | | OTU39 | *Anopheles albimanus* |
| | | | | OTU66 | *Wyeomyia luteoventralis* |
| | | | | AA | *Aedes albopictus* |
| | | OP03B | 2 | OTU01 | *Aedes angustivittatus* |
| | | | | OTU07 | *Limatus durhamii* |
| | | | | OTU40 | *Wyeomyia undulata* |
| | | | | AA | *Aedes albopictus* |
| Golfito | residential | PV01 | 1 | OTU05 | *Wyeomyia adelpha/guatemala* |
| | | | | OTU25 | *Mansonia titillans* |
| | | | | OTU28 | *Culex erythrothorax* |
| | | | | OTU58 | *Trichoprosopon sp.* |
| | | | | AA | *Aedes albopictus* |
| | agriculture | PV02 | 1 | OTU01 | *Aedes angustivittatus* |
| | | | | OTU07 | *Limatus durhamii* |
| | | | | OTU23 | *Culex quinquefasciatus* |
| | | | | OTU35 | *Trichoprosopon pallidiventer* |
| | | PV04 | 1 | OTU23 | *Culex quinquefasciatus* |
| | forest | PV05 | 1 | OTU15 | *Trichoprosopon digitatum* |

**Table S3. Number of sites in each land use category where each mosquito species was present.**

| Species | Agriculture | Forest | Residential | Total |
|---|---|---|---|---|
| *Aedes albopictus* | 5 | 0 | 13 | 18 |
| *Culex quinquefasciatus* | 4 | 1 | 12 | 17 |
| *Culex nigripalpus* | 4 | 5 | 4 | 13 |
| *Wyeomyia adelpha/guatemala* | 3 | 6 | 3 | 12 |
| *Limatus durhamii* | 3 | 0 | 5 | 9 |
| *Wyeomyia complosa* | 2 | 5 | 1 | 8 |
| *Limatus asulleptus* | 2 | 2 | 3 | 7 |
| *Aedes angustivittatus* | 4 | 2 | 0 | 6 |
| *Trichoprosopon digitatum* | 4 | 2 | 0 | 6 |
| *Aedes allotecnon* | 1 | 4 | 0 | 5 |
| *Culex lactator* | 0 | 0 | 5 | 5 |
| *Culex sp. 1* | 2 | 2 | 1 | 5 |
| *Psorophora ferox* | 3 | 2 | 0 | 5 |
| *Trichoprosopon pallidiventer* | 2 | 3 | 0 | 5 |
| *Aedes sp. 1* | 1 | 3 | 0 | 4 |
| *Aedes aegypti* | 1 | 1 | 1 | 3 |
| *Culex coronator* | 1 | 1 | 1 | 3 |
| *Culex erythrothorax* | 1 | 1 | 1 | 3 |
| *Haemagogus sp.* | 2 | 1 | 0 | 3 |
| *Trichoprosopon sp. 3 DJS-2020* | 0 | 3 | 0 | 3 |
| *Wyeomyia sp. RCH-1* | 3 | 0 | 0 | 3 |
| *Coquillettidia nigricans* | 0 | 0 | 2 | 2 |
| *Culex mollis* | 0 | 1 | 1 | 2 |
| *Culex restrictor* | 0 | 2 | 0 | 2 |
| *Haemagogus janthinomys* | 1 | 1 | 0 | 2 |
| *Mansonia titillans* | 1 | 0 | 1 | 2 |
| *Orthopodomyia sp.* | 0 | 1 | 1 | 2 |
| *Trichoprosopon sp. 1* | 0 | 1 | 1 | 2 |
| *unidentified Culicidae 4* | 1 | 1 | 0 | 2 |
| *Wyeomyia luteoventralis* | 1 | 1 | 0 | 2 |
| *Wyeomyia undulata* | 0 | 0 | 1 | 2 |
| *Aedes ferox* | 0 | 1 | 0 | 1 |
| *Aedes infirmatus* | 1 | 0 | 0 | 1 |
| *Aedes sexlineatus* | 0 | 1 | 0 | 1 |
| *Aedes trivittatus* | 0 | 1 | 0 | 1 |

| | | | | |
|---|---|---|---|---|
| *Anopheles albimanus* | 1 | 0 | 0 | 1 |
| *Culex interrogator* | 0 | 0 | 1 | 1 |
| *Culex sp. 2* | 0 | 1 | 0 | 1 |
| *Orthopodomyia MBI-02* | 0 | 0 | 1 | 1 |
| *Psorophora albipes* | 0 | 1 | 0 | 1 |
| *Psorophora funiculus* | 1 | 0 | 0 | 1 |
| *Sabethes undosus* | 0 | 1 | 0 | 1 |
| *Trichoprosopon sp. 5 DJS-2020* | 0 | 1 | 0 | 1 |
| *unidentified Culicidae 1* | 0 | 1 | 0 | 1 |
| *unidentified Culicidae 2* | 0 | 1 | 0 | 1 |
| *unidentified Culicidae 3* | 1 | 0 | 0 | 1 |
| *Wyeomyia aporonoma* | 0 | 0 | 1 | 1 |
| *Wyeomyia sp. 2* | 1 | 0 | 0 | 1 |

**Table S4. Results of generalized linear models estimating the effects of tree cover across spatial scales on species richness. Significant models are highlighted with bold text.**

| Tree cover radius (m) | Estimated effect | Standard error | Z-value | p-value | AIC |
|---:|---:|---:|---:|---:|---:|
| 30 | $3.67 \times 10^{-3}$ | $3.57 \times 10^{-3}$ | 1.03 | $3.03 \times 10^{-1}$ | 194.88 |
| 40 | $5.13 \times 10^{-3}$ | $3.72 \times 10^{-3}$ | 1.38 | $1.77 \times 10^{-1}$ | 194.10 |
| 50 | $5.50 \times 10^{-3}$ | $3.85 \times 10^{-3}$ | 1.43 | $1.54 \times 10^{-1}$ | 193.98 |
| 60 | $6.64 \times 10^{-3}$ | $4.02 \times 10^{-3}$ | 1.65 | $9.8 \times 10^{-2}$ | 193.24 |
| 70 | $7.21 \times 10^{-3}$ | $4.01 \times 10^{-3}$ | 1.80 | $7.23 \times 10^{-2}$ | 192.76 |
| 80 | $8.02 \times 10^{-3}$ | $4.13 \times 10^{-3}$ | 1.94 | $5.24 \times 10^{-2}$ | 192.18 |
| **90** | **$8.93 \times 10^{-3}$** | **$4.25 \times 10^{-3}$** | **2.10** | **$3.58 \times 10^{-2}$** | **191.45** |
| **100** | **$9.13 \times 10^{-3}$** | **$4.30 \times 10^{-3}$** | **2.12** | **$3.39 \times 10^{-2}$** | **191.33** |
| **110** | **$9.64 \times 10^{-3}$** | **$4.37 \times 10^{-3}$** | **2.20** | **$2.75 \times 10^{-2}$** | **190.96** |
| **120** | **$1.05 \times 10^{-2}$** | **$4.46 \times 10^{-3}$** | **2.36** | **$1.84 \times 10^{-2}$** | **190.24** |
| **130** | **$1.06 \times 10^{-2}$** | **$4.47 \times 10^{-3}$** | **2.37** | **$1.76 \times 10^{-2}$** | **190.15** |
| **140** | **$1.10 \times 10^{-2}$** | **$4.58 \times 10^{-3}$** | **2.39** | **$1.66 \times 10^{-2}$** | **190.03** |
| **150** | **$1.17 \times 10^{-2}$** | **$4.62 \times 10^{-3}$** | **2.53** | **$1.16 \times 10^{-2}$** | **189.40** |
| **160** | **$1.20 \times 10^{-2}$** | **$4.67 \times 10^{-3}$** | **2.58** | **$9.92 \times 10^{-3}$** | **189.13** |
| **170** | **$1.24 \times 10^{-2}$** | **$4.69 \times 10^{-3}$** | **2.64** | **$8.18 \times 10^{-3}$** | **188.79** |
| **180** | **$1.29 \times 10^{-2}$** | **$4.77 \times 10^{-3}$** | **2.70** | **$7.01 \times 10^{-3}$** | **188.53** |
| **190** | **$1.30 \times 10^{-2}$** | **$4.83 \times 10^{-3}$** | **2.69** | **$7.09 \times 10^{-3}$** | **188.49** |
| **200** | **$1.35 \times 10^{-2}$** | **$4.87 \times 10^{-3}$** | **2.77** | **$5.59 \times 10^{-3}$** | **188.11** |
| **250** | **$1.40 \times 10^{-2}$** | **$5.00 \times 10^{-3}$** | **2.80** | **$5.08 \times 10^{-3}$** | **187.91** |
| **300** | **$1.37 \times 10^{-2}$** | **$5.01 \times 10^{-3}$** | **2.73** | **$6.27 \times 10^{-3}$** | **188.21** |
| **350** | **$1.38 \times 10^{-2}$** | **$5.09 \times 10^{-3}$** | **2.72** | **$6.57 \times 10^{-3}$** | **188.27** |
| **400** | **$1.37 \times 10^{-2}$** | **$5.24 \times 10^{-3}$** | **2.62** | **$8.79 \times 10^{-3}$** | **188.76** |
| **450** | **$1.33 \times 10^{-2}$** | **$5.36 \times 10^{-3}$** | **2.47** | **$1.34 \times 10^{-2}$** | **189.47** |
| **500** | **$1.26 \times 10^{-2}$** | **$5.48 \times 10^{-3}$** | **2.30** | **$2.17 \times 10^{-2}$** | **190.29** |
| **550** | **$1.19 \times 10^{-2}$** | **$5.54 \times 10^{-3}$** | **2.15** | **$3.13 \times 10^{-2}$** | **190.94** |
| **600** | **$1.15 \times 10^{-2}$** | **$5.58 \times 10^{-3}$** | **2.06** | **$3.93 \times 10^{-2}$** | **191.33** |
| **650** | **$1.11 \times 10^{-2}$** | **$5.63 \times 10^{-3}$** | **1.98** | **$4.80 \times 10^{-2}$** | **191.70** |
| 700 | $1.09 \times 10^{-2}$ | $5.68 \times 10^{-3}$ | 1.92 | $5.51 \times 10^{-2}$ | 191.97 |
| 750 | $1.06 \times 10^{-2}$ | $5.73 \times 10^{-3}$ | 1.85 | $6.43 \times 10^{-2}$ | 192.26 |
| 800 | $1.03 \times 10^{-2}$ | $5.77 \times 10^{-3}$ | 1.78 | $7.44 \times 10^{-2}$ | 192.56 |
| 850 | $1.01 \times 10^{-2}$ | $5.84 \times 10^{-3}$ | 1.73 | $8.38 \times 10^{-2}$ | 192.81 |
| 900 | $9.82 \times 10^{-3}$ | $5.90 \times 10^{-3}$ | 1.66 | $9.64 \times 10^{-2}$ | 193.08 |
| 950 | $9.49 \times 10^{-3}$ | $5.98 \times 10^{-3}$ | 1.59 | $1.13 \times 10^{-1}$ | 193.35 |
| 1000 | $9.23 \times 10^{-3}$ | $6.07 \times 10^{-3}$ | 1.52 | $1.28 \times 10^{-1}$ | 193.57 |

**Table S5. Generalized linear model results estimating effects of percent tree cover calculated at a 1000m radius around survey sites and mean annual temperature on mosquito species richness.**

| Environmental variable | Estimate | Standard Error | Z-value | p-value |
|---|---|---|---|---|
| % tree cover at 1000 m radius | $5.96 \times 10^{-3}$ | $6.24 \times 10^{-3}$ | 0.96 | 0.34 |
| Mean annual temperature | 3.18 | $7.10 \times 10^{-2}$ | $-4.0 \times 10^{-3}$ | 0.99 |
| Tree cover * temperature | $-9.48 \times 10^{-3}$ | $3.48 \times 10^{-3}$ | -2.72 | $6.48 \times 10^{-3}$ * |

**Table S6. Results of generalized linear models estimating the effects of tree cover across spatial scales on *Aedes albopictus* presence. Significant models are highlighted with bold text.**

| Tree cover radius (m) | Estimated effect | Standard error | Z-value | p-value | AIC |
|---|---|---|---|---|---|
| **30** | **$-2.61 \times 10^{-2}$** | **$1.19 \times 10^{-2}$** | **-2.19** | **$2.88 \times 10^{-2}$** | **49.28** |
| **40** | **$-2.95 \times 10^{-2}$** | **$1.31 \times 10^{-2}$** | **-2.25** | **$2.46 \times 10^{-2}$** | **48.70** |
| **50** | **$-3.61 \times 10^{-2}$** | **$1.47 \times 10^{-2}$** | **-2.46** | **$1.39 \times 10^{-2}$** | **46.80** |
| **60** | **$-3.36 \times 10^{-2}$** | **$1.44 \times 10^{-2}$** | **-2.33** | **$1.98 \times 10^{-2}$** | **48.21** |
| **70** | **$-3.51 \times 10^{-2}$** | **$1.48 \times 10^{-2}$** | **-2.37** | **$1.76 \times 10^{-2}$** | **47.79** |
| **80** | **$-4.00 \times 10^{-2}$** | **$1.63 \times 10^{-2}$** | **-2.45** | **$1.44 \times 10^{-2}$** | **46.84** |
| **90** | **$-4.15 \times 10^{-2}$** | **$1.69 \times 10^{-2}$** | **-2.46** | **$1.40 \times 10^{-2}$** | **46.85** |
| **100** | **$-4.09 \times 10^{-2}$** | **$1.72 \times 10^{-2}$** | **-2.38** | **$1.72 \times 10^{-2}$** | **47.32** |
| **110** | **$-4.46 \times 10^{-2}$** | **$1.83 \times 10^{-2}$** | **-2.44** | **$1.47 \times 10^{-2}$** | **46.60** |
| **120** | **$-4.56 \times 10^{-2}$** | **$1.88 \times 10^{-2}$** | **-2.43** | **$1.52 \times 10^{-2}$** | **46.74** |
| **130** | **$-4.52 \times 10^{-2}$** | **$1.88 \times 10^{-2}$** | **-2.41** | **$1.61 \times 10^{-2}$** | **46.91** |
| **140** | **$-4.68 \times 10^{-2}$** | **$1.96 \times 10^{-2}$** | **-2.39** | **$1.69 \times 10^{-2}$** | **46.90** |
| **150** | **$-4.63 \times 10^{-2}$** | **$1.95 \times 10^{-2}$** | **-2.37** | **$1.79 \times 10^{-2}$** | **47.25** |
| **160** | **$-4.71 \times 10^{-2}$** | **$2.00 \times 10^{-2}$** | **-2.36** | **$1.83 \times 10^{-2}$** | **47.28** |
| **170** | **$-4.73 \times 10^{-2}$** | **$2.03 \times 10^{-2}$** | **-2.33** | **$1.96 \times 10^{-2}$** | **47.43** |
| **180** | **$-4.57 \times 10^{-2}$** | **$2.02 \times 10^{-2}$** | **-2.27** | **$2.34 \times 10^{-2}$** | **48.09** |
| **190** | **$-4.52 \times 10^{-2}$** | **$2.03 \times 10^{-2}$** | **-2.23** | **$2.60 \times 10^{-2}$** | **48.40** |
| **200** | **$-4.62 \times 10^{-2}$** | **$2.08 \times 10^{-2}$** | **-2.22** | **$2.63 \times 10^{-2}$** | **48.40** |
| **250** | **$-4.36 \times 10^{-2}$** | **$2.10 \times 10^{-2}$** | **-2.08** | **$3.75 \times 10^{-2}$** | **49.41** |
| 300 | $-3.90 \times 10^{-2}$ | $1.99 \times 10^{-2}$ | -1.96 | $5.00 \times 10^{-2}$ | 50.30 |
| 350 | $-3.67 \times 10^{-2}$ | $1.95 \times 10^{-2}$ | -1.89 | $5.94 \times 10^{-2}$ | 50.82 |
| 400 | $-3.52 \times 10^{-2}$ | $1.92 \times 10^{-2}$ | -1.83 | $6.71 \times 10^{-2}$ | 51.18 |
| 450 | $-3.40 \times 10^{-2}$ | $1.89 \times 10^{-2}$ | -1.80 | $7.24 \times 10^{-2}$ | 51.41 |
| 500 | $-3.35 \times 10^{-2}$ | $1.87 \times 10^{-2}$ | -1.79 | $7.29 \times 10^{-2}$ | 51.48 |
| 550 | $-3.25 \times 10^{-2}$ | $1.84 \times 10^{-2}$ | -1.77 | $7.69 \times 10^{-2}$ | 51.64 |
| 600 | $-3.01 \times 10^{-2}$ | $1.79 \times 10^{-2}$ | -1.68 | $9.25 \times 10^{-2}$ | 52.06 |
| 650 | $-2.77 \times 10^{-2}$ | $1.75 \times 10^{-2}$ | -1.58 | $1.13 \times 10^{-1}$ | 52.49 |
| 700 | $-2.52 \times 10^{-2}$ | $1.72 \times 10^{-2}$ | -1.47 | $1.43 \times 10^{-1}$ | 52.94 |
| 750 | $-2.26 \times 10^{-2}$ | $1.69 \times 10^{-2}$ | -1.34 | $1.81 \times 10^{-1}$ | 53.36 |
| 800 | $-2.10 \times 10^{-2}$ | $1.68 \times 10^{-2}$ | -1.25 | $2.12 \times 10^{-1}$ | 53.62 |
| 850 | $-1.96 \times 10^{-2}$ | $1.67 \times 10^{-2}$ | -1.17 | $2.41 \times 10^{-1}$ | 53.83 |
| 900 | $-1.85 \times 10^{-2}$ | $1.67 \times 10^{-2}$ | -1.10 | $2.70 \times 10^{-1}$ | 54.00 |
| 950 | $-1.75 \times 10^{-2}$ | $1.68 \times 10^{-2}$ | -1.04 | $2.98 \times 10^{-1}$ | 54.15 |
| 1000 | $-1.65 \times 10^{-2}$ | $1.69 \times 10^{-2}$ | -0.98 | $3.28 \times 10^{-1}$ | 54.28 |

**Table S7. Generalized linear model results estimating effects of percent tree cover calculated at a 1000m radius around survey sites and mean annual temperature on *Ae. albopictus* presence.**

| Environmental variable | Estimate | Standard Error | Z-value | p-value |
|---|---|---|---|---|
| % tree cover at 1000 m radius | $-8.34 \times 10^{-2}$ | $3.61 \times 10^{-2}$ | -2.31 | 0.021 * |
| Mean annual temperature | $9.37 \times 10^{-1}$ | $4.20 \times 10^{-2}$ | 2.23 | 0.026 * |
| Tree cover * temperature | $-3.46 \times 10^{-3}$ | $1.54 \times 10^{-2}$ | -0.225 | 0.822 |

**Table S8. Generalized dissimilarity modeling results for mosquito community turnover along environmental gradients of mean annual temperature, elevation, geographic distance, and tree cover calculated for different radii surrounding survey sites.**

| | *sum of coefficients* | | | | | |
|---|---|---|---|---|---|---|
| **Tree cover radius (m)** | **percent tree cover** | **mean annual temperature** | **geographic distance** | **elevation** | **intercept** | **percent deviance explained** |
| 30 | 0.659 | 0.916 | 0.410 | 0 | 1.64 | 5.430 |
| 40 | 0.602 | 0.890 | 0.417 | 0 | 1.64 | 5.727 |
| 50 | 0.725 | 0.858 | 0.394 | 0 | 1.60 | 6.797 |
| 60 | 0.698 | 0.904 | 0.420 | 0 | 1.63 | 6.046 |
| 70 | 0.694 | 0.900 | 0.408 | 0 | 1.63 | 6.036 |
| 80 | 0.750 | 0.875 | 0.388 | 0 | 1.61 | 6.472 |
| 90 | 0.737 | 0.880 | 0.409 | 0 | 1.61 | 6.367 |
| 100 | 0.710 | 0.889 | 0.399 | 0 | 1.62 | 6.110 |
| 110 | 0.788 | 0.873 | 0.372 | 0 | 1.59 | 6.651 |
| 120 | 0.801 | 0.878 | 0.376 | 0 | 1.59 | 6.643 |
| 130 | 0.823 | 0.875 | 0.356 | 0 | 1.59 | 6.803 |
| 140 | 0.840 | 0.873 | 0.349 | 0 | 1.59 | 6.745 |
| 150 | 0.831 | 0.872 | 0.347 | 0 | 1.59 | 6.590 |
| 160 | 0.850 | 0.868 | 0.333 | 0 | 1.59 | 6.566 |
| 170 | 0.837 | 0.865 | 0.322 | 0 | 1.60 | 6.393 |
| 180 | 0.799 | 0.860 | 0.340 | 0 | 1.62 | 5.821 |
| 190 | 0.773 | 0.858 | 0.357 | 0 | 1.65 | 5.472 |
| 200 | 0.792 | 0.850 | 0.356 | 0 | 1.65 | 5.422 |
| 250 | 0.628 | 0.856 | 0.455 | 0 | 1.74 | 4.383 |
| 300 | 0.520 | 0.880 | 0.520 | 0 | 1.79 | 4.106 |
| 350 | 0.499 | 0.884 | 0.527 | 0 | 1.79 | 4.094 |
| 400 | 0.465 | 0.889 | 0.513 | 0 | 1.79 | 4.095 |
| 450 | 0.434 | 0.905 | 0.515 | 0 | 1.79 | 3.947 |
| 500 | 0.405 | 0.921 | 0.517 | 0 | 1.79 | 3.812 |
| 550 | 0.372 | 0.937 | 0.525 | 0 | 1.80 | 3.690 |
| 600 | 0.314 | 0.543 | 0.950 | 0 | 1.80 | 3.544 |
| 650 | 0.258 | 0.558 | 0.962 | 0 | 1.81 | 3.424 |
| 700 | 0.211 | 0.571 | 0.973 | 0 | 1.81 | 3.339 |
| 750 | 0.163 | 0.583 | 0.982 | 0 | 1.82 | 3.272 |
| 800 | 0.119 | 0.593 | 0.990 | 0 | 1.82 | 3.224 |
| 850 | 0.066 | 0.608 | 0.999 | 0 | 1.83 | 3.184 |
| 900 | 0.023 | 0.623 | 1.004 | 0 | 1.83 | 3.166 |
| 950 | 0.001 | 0.631 | 1.006 | 0 | 1.84 | 3.163 |
| 1000 | 0 | 0.631 | 1.007 | 0 | 1.84 | 3.163 |
